# Progenitor cell integration into a barrier epithelium during adult organ turnover

**DOI:** 10.1101/2021.09.19.457819

**Authors:** Paola Moreno-Roman, Yu-Han Su, Anthony Galenza, Lehi Acosta-Alvarez, Alain Debec, Antoine Guichet, Jon-Michael Knapp, Caroline Kizilyaprak, Bruno M. Humbel, Irina Kolotuev, Lucy Erin O’Brien

## Abstract

Barrier epithelial organs face the constant challenge of sealing the interior body from the external environment while simultaneously replacing the cells that contact this environment. These replacement cells—the progeny of basal stem cells—are born without apical, barrier-forming structures such as a protective, lumen-facing membrane and occluding junctions. How stem cell progeny acquire these structures to become part of the barrier is unknown. Here we use Focused Ion Beam-Scanning Electron Microscopy (FIB-SEM), Correlative Light-Electron Microscopy (CLEM), and volumetric imaging of live and fixed organs to investigate progenitor integration in the intestinal epithelium of adult *Drosophila*. We find that stem cell daughters gestate their future lumenal-apical membrane beneath a transient, basal niche formed by an umbrella-shaped occluding junction that shelters the growing cell and adheres it to mature neighbor cells. The umbrella junction both targets formation of a deep, microvilli-lined, apical invagination and closes it off from the contents of the gut lumen. When the growing cell is sufficiently mature, the umbrella junction retracts to expose this Pre-Assembled Apical Compartment (PAAC) to the gut lumen, thus incorporating the new cell into the intestinal barrier. When we block umbrella junctions, stem cell daughters grow and attempt to differentiate but fail to integrate; when we block cell growth, no umbrella junctions form and daughters arrest in early differentiation. Thus, stem cell progeny build new barrier structures in the shelter of a transient niche, where they are protected from lumenal insults until they are prepared to withstand them. By coordinating this dynamic junctional niche with progenitor cell differentiation, a physiologically active epithelial organ incorporates new cells while upholding integrity of its barrier.

## INTRODUCTION

Barrier epithelial organs protect the body interior from the external environment while performing physiological processes that require direct exposure to this environment. For example, the epithelium of the digestive tract both protects the body from gastric acid and enteric pathogens while simultaneously breaking down and absorbing ingested nutrients. The dual roles of barrier epithelia create a conundrum: Optimal physiological function requires that the tissue replace old, spent cells with new stem cell progeny, but each individual progeny must be incorporated into the epithelium without compromising barrier integrity (Guillot and Lecuit, 2013; Leblond, 1981; Liang et al., 2017; Macara et al., 2014; Pellettieri and Alvarado, 2007). The cellular mechanisms which enable stem cell progeny to seamlessly assimilate into a functioning barrier are poorly understood.

In all metazoans, epithelial barrier function arises from two conserved features of the epithelial cells themselves. First, cell-cell occluding junctions—tight junctions in vertebrates, septate junctions in invertebrates—create impermeable seals between cells (Varadarajan et al., 2019). Occluding junctions encircle each epithelial cell at the lateral border of the apical membrane, creating a sealed network that prevents even small molecules from freely passing between the lumen and the body interior. Second, a lumen-facing, apical plasma membrane, tightly folded into microvilli or cilia, forms a mucosal shield that resists corrosives, pathogens, and other lumenal insults (Linden et al., 2008; McGuckin et al., 2011; Overeem et al., 2015). This apical membrane is a hallmark of epithelial differentiation and serves as the barrier’s direct interface with the outside world. In contrast, the stem cells that renew many barrier epithelia lack occluding junctions and a lumen-contacting apical membrane. Examples of such epithelia include the mammalian trachea (Evans and Moller, 1991; Michael J. Evans, 2001; Rock et al., 2009; Sekiya et al., 1988), mammary gland (Chepko and Dickson, 2003; Chepko and Smith, 1997), prostate (Tsujimura et al., 2002), cornea (Cotsarelis et al., 1989), and olfactory lining (Leung et al., 2007), and the *Drosophila* adult midgut (Korzelius et al., 2014; Resnik-Docampo et al., 2017; Xu et al., 2019). Stem cells in these tissues are much smaller than their mature progeny; they inhabit the basal region of the epithelium, protected from lumenal contents by the mature cells’ occluding junction network.

Since stem cells lack barrier-forming structures, their progeny must generate these structures *de novo* as they integrate into the barrier during terminal differentiation. Radial intercalation has been proposed to be this integration mechanism (Walck-Shannon and Hardin, 2014; Sedzinski et al., 2016; Chen et al., 2018). Many developing epithelial tissues use radial intercalation to merge basally derived cells into an overlying epithelium (Merzdorf et al., 1998; Deblandre et al., 1999; Stubbs et al., 2006; Voiculescu et al., 2007; McMahon et al., 2008; Campbell et al., 2010). In this process, a cell that is born basal to the occluding junction network integrates into this network by moving apically while wedging itself between pre-existing cells. When the tip of the intercalating cell reaches the epithelium’s occluding junctions, the new cell forms occluding junctions with its neighbors. These SJs begin as a pinpoint and morph into a ring that surrounds the cell’s nascent, lumen-facing apical membrane. This apical membrane and its encircling occluding junction expand radially as the cell grows to its final size (Stubbs et al., 2006; Sedzinski et al., 2016, 2017). Similar to these developmental contexts, integration of adult stem cell progeny involves basal-to-apical movement and *de novo* formation of barrier-forming structures. Whether stem cell progeny use radial intercalation or an alternate, perhaps novel, mechanism, remains unexamined.

We leveraged recent advances in Focused Ion Beam-Scanning Electron Microscopy (FIB-SEM), Correlative Light-Electron Microscopy (CLEM – (Burel et al., 2018)), and *in vivo* volumetric confocal live imaging (Martin et al., 2018) to directly examine this question using the midgut of adult *Drosophila*. Like many vertebrate barrier epithelia, the epithelial lining of the fly midgut is a leakproof, pseudostratified epithelium that is continually renewed through the divisions of basal stem cells (Lemaitre and Miguel-Aliaga, 2013). Investigating how new stem cell progeny assimilate into this adult barrier epithelium, we found—unexpectedly—that assimilation occurs not by radial intercalation but by a striking morphogenetic process that has not previously been described for any tissue.

We discovered that as a stem cell daughter undergoes terminal differentiation, its nascent occluding junctions form a transient, umbrella-shaped niche that supports development of the cell’s future, lumen-facing apical surface. This surface starts as an intercellular, localized delamination within the umbrella-shaped junction. The delaminated membrane of the differentiating cell accumulates apical markers, invaginates deeply into the cell’s cytoplasm, and folds into microvilli in the shelter of the junctional niche; these behaviors create a <u>p</u>re-<u>a</u>ssembled <u>a</u>pical <u>c</u>ompartment (PAAC) on the basal side of the epithelial barrier, protected from the contents of the gut lumen. In the final phase of cell differentiation, the umbrella junction retracts to fuse the PAAC with the gut lumen, and the apical membrane everts to form the mature cell’s convex lumenal surface.

The morphogenetic process of progenitor cell integration is coupled to growth that progenitor cells undergo as they terminally differentiate. When we block stem cell daughters from integrating, they become trapped in a hybrid, partially differentiated state and accumulate on the basal side of the epithelium. These animals die prematurely, implying that organismal longevity is compromised when the intestinal barrier is not properly replenished.

We suggest that PAAC-mediated integration enables stem cell progeny to generate lumen-facing cell surfaces in a space that is protected from lumenal insults, thus enabling new cells to be added seamlessly to a physiologically active barrier epithelium.

## RESULTS

Mature intestinal enterocytes form the bulk of the *Drosophila* midgut and are responsible for its barrier function. Like their vertebrate counterparts, *Drosophila* enterocytes are bonded together by apical occluding junctions (Figs. 1A & 2A). In the fly gut, these occluding junctions are smooth septate junctions (SJs) (Furuse and Izumi, 2017). Also like their vertebrate counterparts, *Drosophila* enterocytes display an apical brush border composed of long, dense microvilli (Fig. 2A); the brush border both absorbs nutrients and protects against lumenal pathogens.

**Figure 1.**
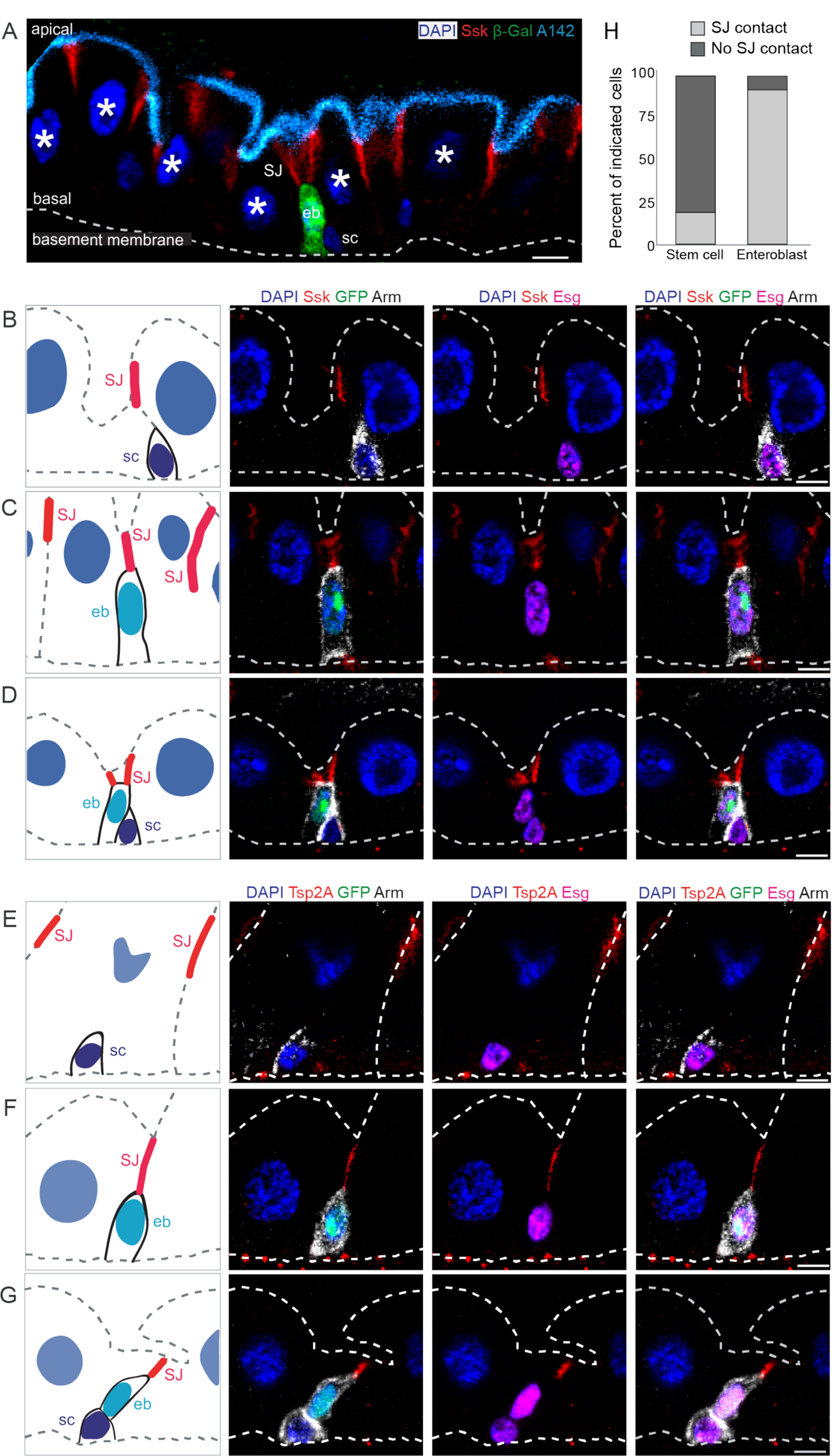
The apical tip of a differentiating enteroblast contacts the SJ of its neighbor enterocytes. (A) Architecture and stem cell lineage of the fly midgut epithelium, shown in cross-sectional view with apical lumenal surface (cyan, Mdu::GFP; *c.f.* Fig. S2) at top and basal surface (dotted line, basement membrane) at bottom. Three cell types make up the absorptive lineage: (1) Stem cells (sc) are basally localized, diploid cells that do not express *Su(H)-lacZ*. (2) Enteroblasts (eb) are terminally committed stem cell progeny. Enteroblasts are transitioning from stem-like cells to enterocytes and are marked by *Su(H)-lacZ* expression (green, β-Gal). Stem cells and enteroblasts often appear in pairs. (3) Mature enterocytes are large cells with polyploid nuclei (asterisks). Septate junctions (SJ; red, Snakeskin) appear at the apico-lateral borders of enterocytes. (B-G) Stem cells do not overlap with SJs, while the apical tips of enteroblasts contact the basal termini of enterocyte-enterocyte SJs. Cartoons (left column) and channel overlays from 5-channel multi-photon laser microscopy of *esgGAL4, UAS-his2b::CFP; Su(H)-GFP:nls* midguts immunostained for SJ components Ssk (red, B-D) or Tsp2a (red, E-G) and for the stem cell/enteroblast marker Arm (white; cortical). *esg*-driven His2b::CFP is shown in magenta, *Su(H)*-driven GFP:nls in green and nuclei (DAPI) in blue. Lumenal epithelial surface and basement membrane are indicated by dotted gray lines. Stem cells (sc) are His::CFP^+^, Arm^+^, GFP:nls^-^ cells in Panels B, D, E, G; enteroblasts (eb) are His::CFP^+^, Arm^+^,GFP:nls^+^ cells in Panels C, D, F, G. Panels D and G show stem cell-enteroblast pairs. All scale bars: 5µm. Images are projections of short confocal stacks. Full genotypes in Table 1. (H) Quantitation of B-G. Most enteroblasts (92%), but few stem cells (19%) contact the epithelial septate junction network. N=5 midguts; n=119 stem cells and 125 enteroblasts.

**Figure 2.**
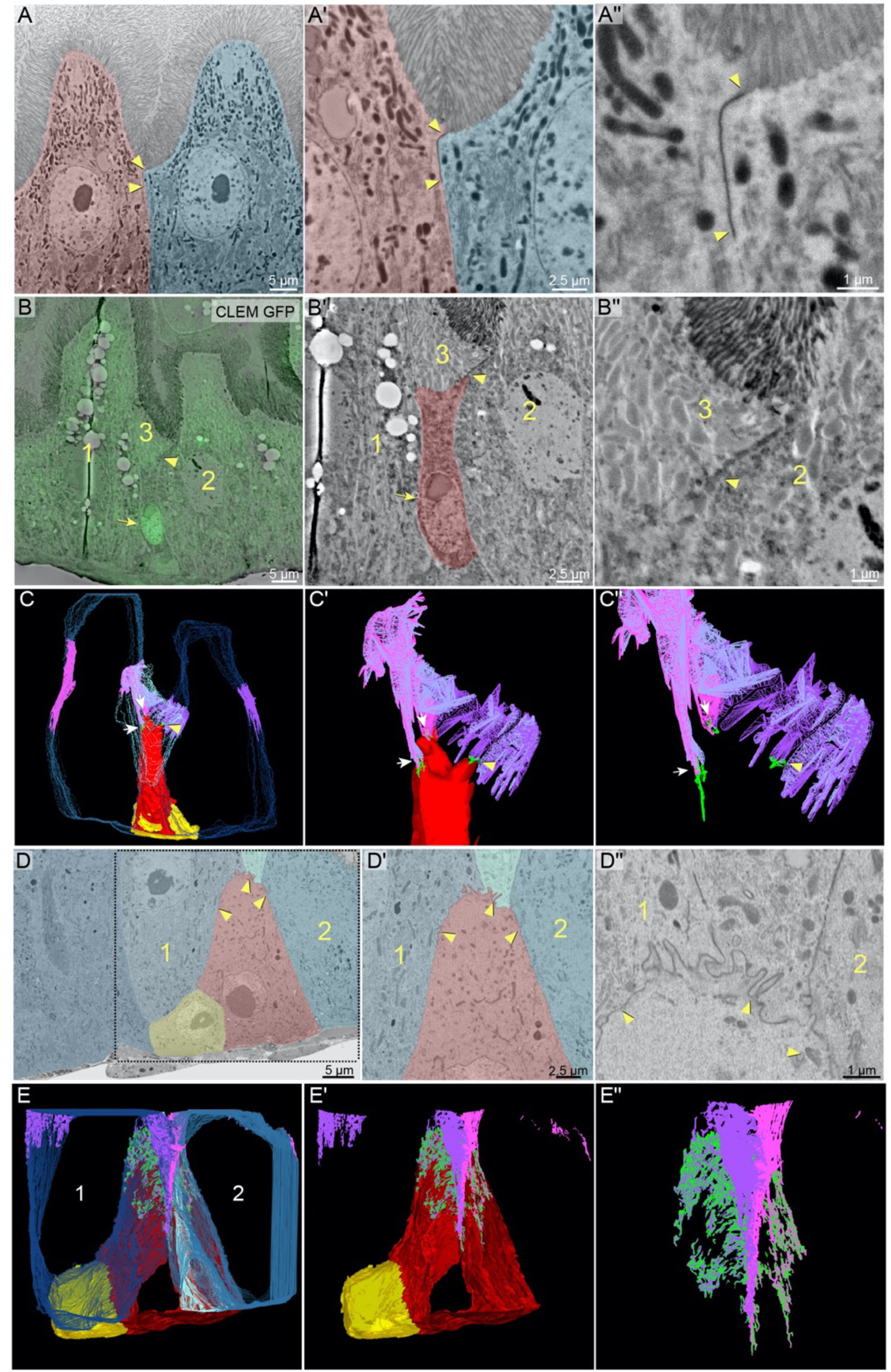
The differentiating enteroblast’s apex initiates a new SJ at the basal-most edge of mature enterocyte SJs and triggers their remodeling to form a basally-extended SJ sheet. (A) Mature enterocytes localize SJs to the boundary between lumen-facing, brush border apical membranes and lateral membranes. SEM shows two mature enterocytes (pink and blue pseudocolor). Apical membranes are identifiable as microvilli-rich brush borders. An electron-dense SJ (arrowheads) fuses together the enterocytes’ lateral membranes at a site directly adjacent to these brush borders. Zoomed-in views of SJ in A are shown in A’ and A’’. (B-C) The apex (apical-most tip) of a young, Su(H)^+^ enteroblast initiates SJ adhesions at the basal edge of enterocyte SJs. CLEM overlay (B) identifies a Su(H)-GFP::nls^+^ enteroblast (arrow) in a FIB-SEM section. Zoomed-in images of the enteroblast (B’, red pseudocolor) and of the enteroblast apex (B’’) show a nascent SJ (arrowheads in B-B’’) between the enteroblast and neighbor enterocytes 2 and 3 (labelled). (Only a small wedge of enterocyte 3 is visible in this section.) See Fig. S1. Volumetric rendering (C) of 30 FIB-SEM sections, including the section in B, reveals that each of apex’s three fingers forms a SJ (arrows and arrowhead; arrowhead points to the same SJ in B and C) with each of three neighbor enterocytes. Cells and SJs are color coded: enteroblast, red; enteroblast SJs, green; enterocytes 1-3, blue; enterocyte 1 SJs, magenta; enterocyte 2 SJs, light purple; enterocyte 3 SJs, light blue; stem cell, yellow. Zoomed-in views of the enteroblast apex and associated SJs are shown in C’ and C’’. See Video 1. (D-E) An older enteroblast is blanketed by the broad, basally extended SJ it has formed with the lateral membranes of neighboring mature cells. A FIB-SEM section (D) shows an enteroblast (red pseudocolor), two mature enterocytes (cells 1 and 2; blue pseudocolor), a mature enteroendocrine cell (cell 3; light blue pseudocolor), and a presumptive stem cell (yellow pseudocolor). The apical third of the enteroblast has formed an SJ (arrowheads in D-D’’) with the lateral membranes of the mature cells. Zoomed-in views of the enteroblast apex are shown in D’ and D’’. Volumetric rendering (E) of 413 FIB-SEM sections, including the section in D, reveals basal extensions of both enteroblast-enterocyte SJs and associated enterocyte-enterocyte SJs. View of the SJ with only the enteroblast and stem cell is shown in E’. Zoomed-in view of the SJ alone is shown in E’’. Cells and SJs are color coded: enteroblast, red; enteroblast SJ, green; enterocytes 1 and 2, blue; enterocyte 1 SJ, lavender; enterocyte 2 SJ, magenta; enteroendocrine cell, light blue; stem cell, yellow. See Video 2. Full genotypes in Table 1.

**TABLE 1.**
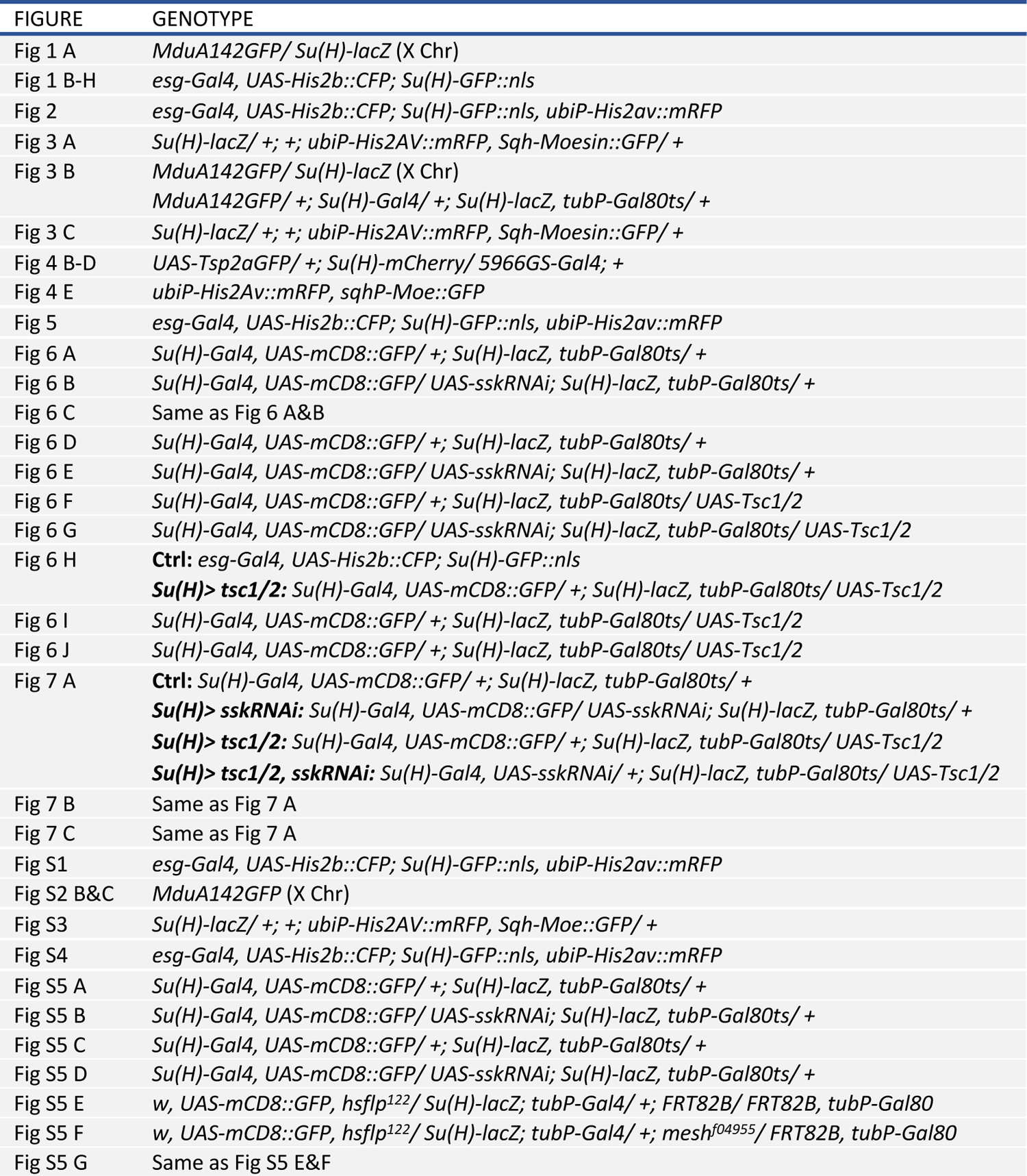
Genotypes in Figure Panels

**TABLE 2.**
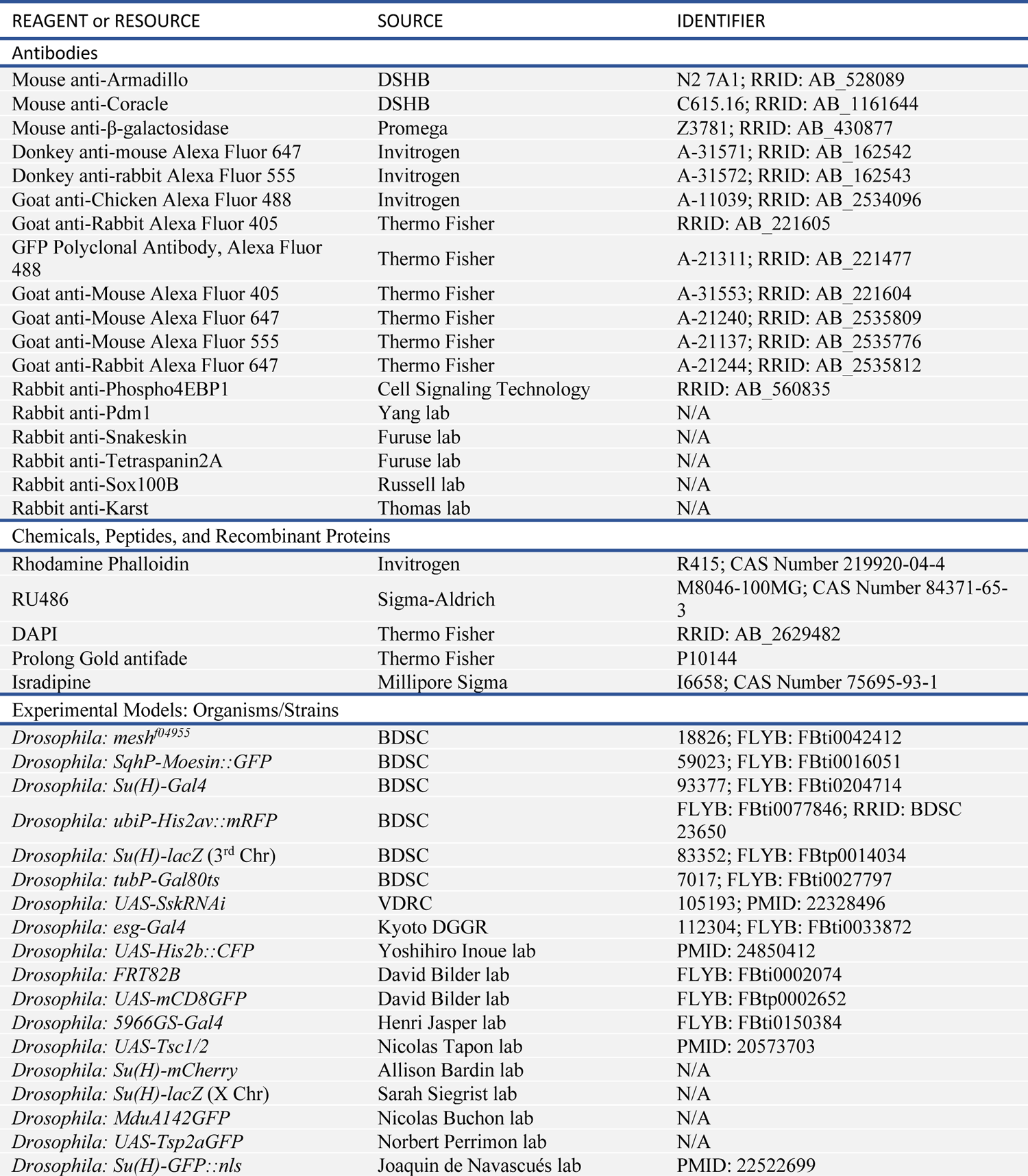
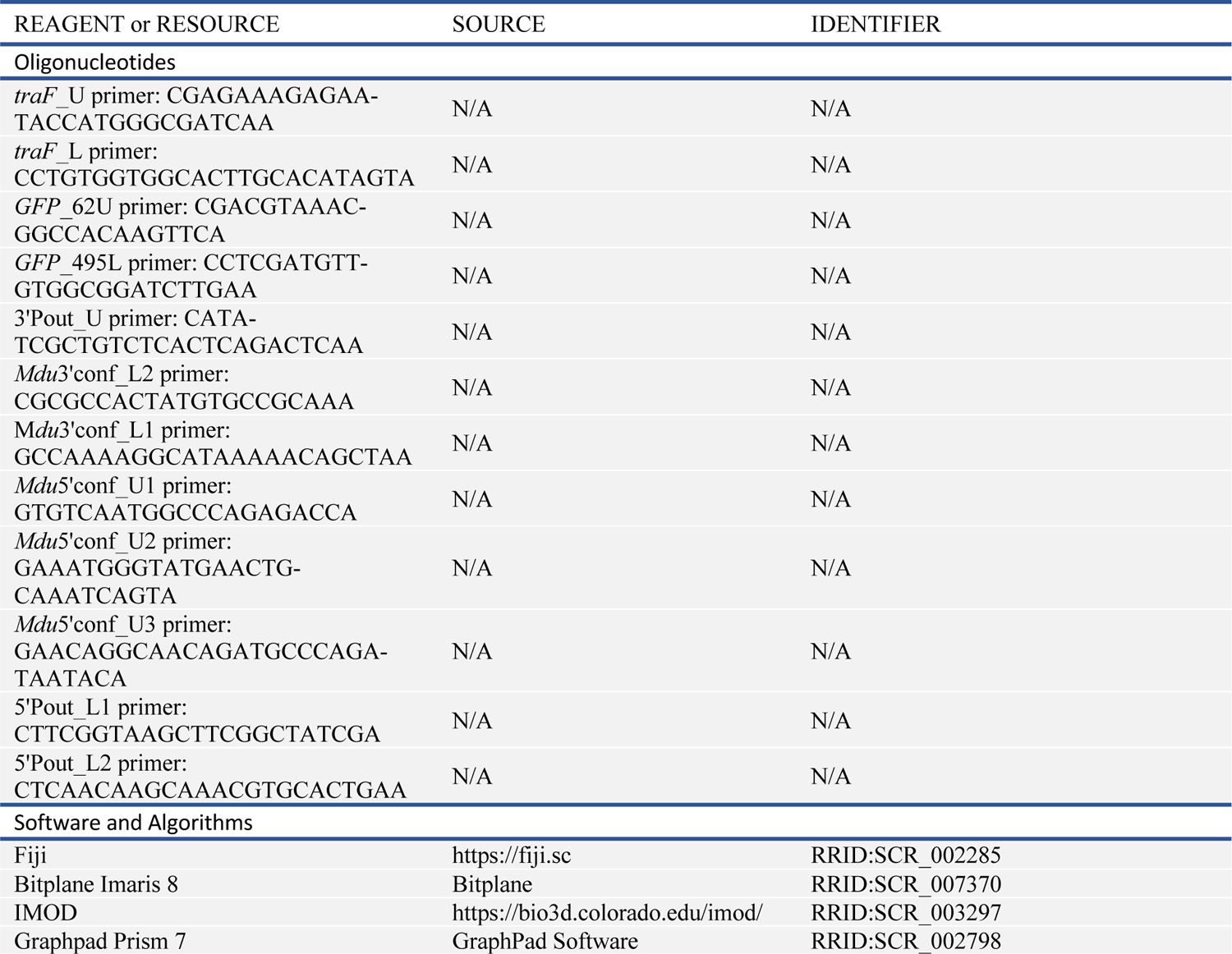
Reagents and Resources

Enterocytes are terminally differentiated and post-mitotic. When shed through damage or death, they are replaced by division of resident stem cells (Micchelli and Perrimon, 2006; Ohlstein and Spradling, 2006; Jiang and Edgar, 2009; Liang et al., 2017). Stem cell progeny that are fated to become enterocytes must first pass through an intermediate, post-mitotic stage called an enteroblast (Ohlstein and Spradling, 2007; Bardin et al., 2010; Perdigoto et al., 2011) (Figure 1A). Enteroblast identity is determined by activation of the Notch receptor and can be visualized using Notch reporters such as *Su(H)-GFP::nls* and *Su(H)-lacZ* (Ohlstein and Spradling, 2007; Bardin et al., 2010; Perdigoto et al., 2011; de Navascués et al., 2012). As enteroblasts differentiate to enterocytes, they endoreplicate from 2N to 32-64N and increase in volume by ∼30x (Xiang et al., 2017).

### Stem cell progeny initiate new SJs at discrete contact points with mature cells

We were intrigued by prior observations that stem cells lack SJs (Korzelius et al., 2014; Resnik-Docampo et al., 2017; Xu et al., 2019) because this implies that differentiating cells must form SJs *de novo* and integrate themselves into the enterocytes’ leakproof SJ network. We sought to determine when, where, and how *de novo* SJ formation occurs. First, we asked whether enteroblasts form SJs with their mature enterocyte neighbors (Figs. 1 & 2). If so, then SJ components should localize to enteroblast-enterocyte interfaces. To investigate this prediction, we used midguts that expressed markers to distinguish enteroblasts, enterocytes, and stem cells. We immunostained them for the SJ components Snakeskin (Ssk) and Tetraspanin2A (Tsp2A) (Yanagihashi et al., 2012; Izumi et al., 2016), then performed 5-channel multi-photon laser scanning microscopy to visualize SJs in the context of identified cells.

We observed SJ components at nearly all enteroblast-enterocyte interfaces, but not at the vast majority of stem cell-enterocyte interfaces nor at any stem cell-enteroblast interfaces. 89% of stem cells (*escargotGAL4, UAS-his2b::CFP* (*esg*^+^), Armadillo (Arm^+^), Su(H)-GFP:nls^-^) exhibited no co-localization with SJ components (n=119 stem cells from 5 guts) (Fig. 1B & 1E), as expected (Korzelius et al., 2014; Resnik-Docampo et al., 2017; Xu et al., 2019). By contrast, 92% of enteroblasts (*esg*^+^, Arm^+^, Su(H)-GFP:nls^+^) (Figs. 1C, 1D, 1F & 1G) overlapped with enterocyte-enterocyte SJs (n=125 enteroblasts from 5 guts). Many SJ-contacting enteroblasts exhibited small, presumably diploid, nuclei, which suggests that these contacts are formed in initial stages of enteroblast fate determination.

Enteroblast SJ staining invariably localized to the apical-most tips (apex) of the enteroblasts, where it overlapped with the basal terminus of the enterocyte SJs. This overlap might represent either *bona fide* adhesion septa between enteroblasts and enterocytes or else the mere physical proximity without formation of true adhesion septa. To distinguish these scenarios, we performed correlative light-electron microscopy (CLEM) (Kolotuev, 2014) on *Su(H)-GFP::nls*-expressing midguts to identify the GFP-labelled enteroblasts. FIB was performed in random positions of the midgut R4c region; GFP-labeled enteroblasts were identified; and interfaces between these enteroblasts and neighbor enterocytes were examined for potential SJ septa.

In EM, SJ septa characteristically appear as electron-dense structures that ‘fuse’ together the apposing plasma membranes of adjacent cells. As expected, enterocyte-enterocyte septa fused together the apical-most regions of the cells’ lateral membranes, directly adjacent to their microvilli-rich brush borders (Fig. 2A). However, when enterocytes were next to an enteroblast (Figs. 2B-2E), their SJs extended basally toward the enteroblast (red pseudocolor in Fig. 2B’’) and fused with enteroblast plasma membrane at its apex (arrowheads). These observations demonstrate that enteroblasts form *bona fide* SJ septa with mature neighbor cells. In addition, they imply that light-microscopy visualized overlap of immunostained SJ components with enteroblast cell membranes (Fig. 1) are sites of new SJ formation.

To understand the three-dimensional structure of enteroblast-enterocyte SJs, we performed array tomography. We transformed the tomograms into volumetric renderings of individual cells and their SJ interfaces. Fig. 2C and Video 1 show the volumetric rendering of a 30-slice series surrounding the CLEM image in Fig. 2B. The apex of this diploid (presumably, newly determined) enteroblast (red) exhibits three finger-like projections (arrowheads), each of which forms a point-like, discrete SJ (green) with one of its three neighbor enterocytes (blue; labelled 1-3 in Fig. 2B). No SJs are visible between the enteroblast and an adjacent, Su(H)-GFP:nls-negative stem cell (yellow). The enteroblast’s three point-like SJs contact the three enterocyte SJs (pink, pale blue, and purple) at the latters’ basal rim. Importantly, we found no discontinuities between the new, enteroblast-enterocyte SJs (overlap between green and either pink, pale blue, or purple) and the pre-existing, enterocyte-enterocyte SJs (overlap between pink and pale blue, pink and purple, or pale blue and purple). This structural continuity suggests that the pre-existing, enterocyte-enterocyte SJ guides the localization of initial SJ contacts between enterocytes and the new enteroblast.

### SJ contacts develop into a large, umbrella-shaped SJ that covers the enteroblast apex and extends toward the basal epithelium

We next generated a volumetric rendering of a larger (hence, likely older) enteroblast (red) from a 413-slice FIB-SEM tomographic series that spanned a 35.6µm x 35.6µm x 4.5µm tissue volume. A single slice from this series is shown in Figs. 2D-D’’ and Fig. S1, and the volumetric rendering is shown in Figure 2E and Video 2. In contrast to the nascent SJ contacts formed by a young enteroblast (Figures 2B & 2C), the SJ (green) of this older enteroblast is an expansive, adhesive ‘zone’ whose shape resembles an umbrella. This umbrella SJ fuses the pyramidal enteroblast to three mature neighbor cells: the SJs (pink and purple) of two enterocytes (blue; labelled 1 and 2 in Figs. 2D & 2E) and those of an enteroendocrine cell (pale blue; labelled 3 in Fig. 2D). No SJs are visible at the interface between the enteroblast and a presumptive stem cell (yellow). As with SJs of the young enteroblast (Fig. 2B-C), the SJs between the older enteroblast and its neighbor enterocytes (green/pink overlap and green/purple overlap) are continuous with the SJ that had previously formed between the enterocytes themselves (pink/purple overlap).

Importantly, the umbrella SJ extends unusually far toward the basal epithelium—a feature that is incompatible with a simple model of radial intercalation. SJs typically occupy a narrow band at the apical border of the enterocytes’ lateral membranes (Fig. 2A; also in Fig. 2E and E’, the purple SJ on left side of enterocyte 1 and pink SJ on right side of enterocyte 2). In radial intercalation, the SJ initiates at the point where the new cell’s apical tip becomes co-planar with the organ’s mature SJ network; hence, the new SJ grows to a narrow width that matches that of mature SJs (Walck-Shannon and Hardin, 2014; Stubbs et al., 2006; Sedzinski et al., 2016, 2017; Chen et al., 2018).

However, an enteroblast’s umbrella SJ spreads basally along two-thirds of the enterocytes’ lateral membranes and shrouds the top third of the enteroblast (Fig. 2E, green SJ and associated pink and purple SJs; Video 2). This basally-extended umbrella shape implies an alternate mode of integration in which the new SJ forms basal to the organ’s mature SJ network.

### Cloning Meduse, an actin-associated protein that localizes to the brush border of midgut enterocytes

The second component of the gut’s barrier structure is collectively formed by the apical cell surfaces that line the gut lumen. Mature enterocytes fold these surfaces into long, dense microvilli, forming the intestinal brush border (Fig. 2A). Mature enterocytes also exhibit apical-basal polarity, as characterized by lumen-polarized localization of cytoskeleton-associated proteins such as Moesin, Karst (β_H_-spectrin), and Myosin7a (Baumann, 2001; Chen et al., 2018).

Another luminally polarized marker is provided by the splice-trap transposon line A142(Bobinnec et al., 2003; Buchon et al., 2013), which expresses a GFP fusion protein that co-localizes with Moesin at or near enterocyte microvilli (Fig. 3A-C). We found that this transposon is inserted into CG2556 (Fig. S2A), a previously uncharacterized gene that does not appear to have vertebrate homologs. Since the filamentous appearance of the A142 fusion protein in egg chambers is reminiscent of sea jelly tentacles (Fig. S2B), we named this gene Meduse (Mdu). Mdu is predicted to be a 470 amino-acid, 51 kDa protein whose sole identifiable motif is an actin binding domain. This putative actin-binding function is consistent with localization of the A142 splice trap to the apical brush border of enterocytes and with actin filaments in Stage 10 egg chambers, the latter of which is latrunculin-sensitive (Fig. S2B-C).

**Figure 3.**
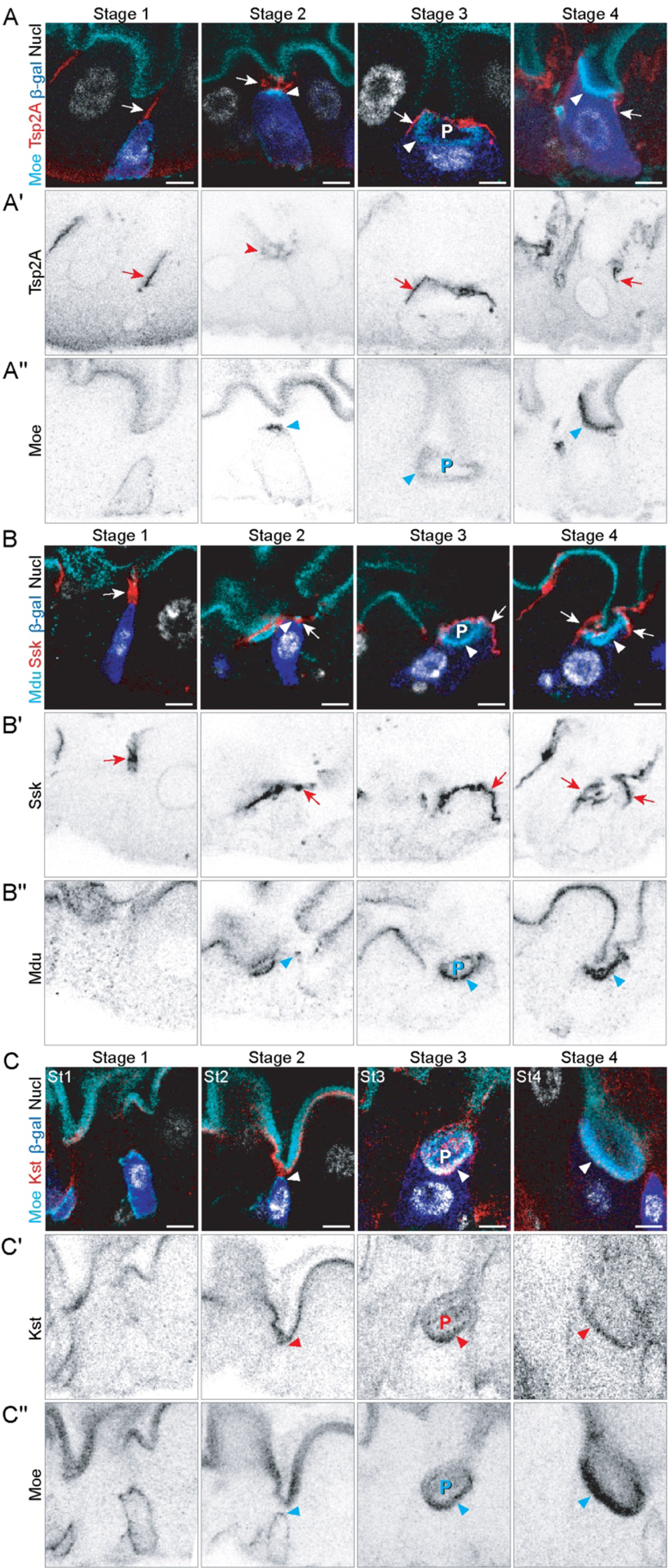
SJ and apical membrane morphology define four stages of barrier integration. As differentiating cells integrate, they pass through four morphological stages that are distinguishable by SJ localization, polarization of apical markers, and cellular/nuclear size. SJs shown in red (A, Tsp2A; B, Ssk). Markers of enterocyte apical polarity shown in either cyan (A, C Moe::GFP; B, Mdu::GFP; *c.f.* Fig. S2) or red (C, Kst). Su(H)-lacZ in blue (β-Gal, A-C). Nuclei (Nucl) shown in grayscale (A, Stage 1 and C – His2av::mRFP; A, Stages 2-4 and B – DAPI). Images are projections of short confocal stacks. Markers that are not fused to a fluorescent protein were visualized by immunostaining. All scale bars, 5 µm. β-gal channel shown in Fig. S3. Full genotypes in Table 1. <u>Stage 1 (St1)</u>. Enteroblast with initial SJ contact. The apex of a diploid enteroblast contacts the basal terminus of an enterocyte-enterocyte SJ (arrows). Apical markers are either non-polarized (Moe, Kst) or absent (Mdu). <u>Stage 2 (St2)</u>. Enteroblast with broadened SJ contacts and apical plaque. The enteroblast apex is partially covered by a widened enteroblast-enterocyte SJ (arrows). Some apical markers (Moe, Mdu; arrowheads) become polarized to the apex, forming a plaque immediately basal to the SJ. <u>Stage 3 (St3)</u>. Pre-enterocyte with umbrella-like SJ and concave apical structure (PAAC). The apex of the cell is fully covered by a convex SJ (arrows). Apical markers outline a prominent concave structure (P; Pre-Assembled Apical Compartment, or PAAC) beneath the SJ. The PAAC fills the apex of the cell and is separate from the gut lumen (Video 3). Pre-enterocytes exhibit cytoplasmic/nuclear sizes intermediate between enteroblasts and mature enterocytes and low *Su(H)*-driven β-gal signal (Fig. S3). <u>Stage 4 (St4)</u>. Pre-enterocyte integration becomes complete. The SJ circumscribes the cell, and the PAAC has coalesced with the gut lumen. The pre-enterocyte is still smaller than mature enterocytes and has a concave, rather than convex apical-lumenal surface, but the relative arrangement of its SJ, apical, and basolateral surfaces are topologically equivalent to a mature enterocyte.

### The umbrella SJ is a transient niche for formation of the new cell’s future lumenal-apical surface

During development of tubular epithelial organs, cells generally couple the formation of a lumen-contacting cell surface to the formation of apical features including microvilli, occluding junctions, and apically polarized membrane and cytoskeletal proteins (Blasky et al., 2015; Datta et al., 2011; O’Brien et al., 2002; Sigurbjörnsdóttir et al., 2014). By comparison, stem cells in the adult fly gut lack this entire suite of lumenal-apical features (Figs. 2B-2E) (Chen et al., 2018). How do stem cell progeny generate a lumenal-apical surface as they differentiate, and how do they coordinate apical morphogenesis with SJ formation and epithelial integration?

To address this question, we used high-resolution images of fixed guts to identify cells at distinct stages of differentiation, assessed the localization of apical markers, and correlated apical marker localization to SJ maturation (Figs. 3 & S2 and Video 3). Midguts that express Su(H)-lacZ were used because the long perdurance of β-galactosidase (>20 h half-life (Bachmair et al., 1986)) makes it possible to identify early-stage enterocytes that have recently turned off enteroblast-specific *Su(H)* activity but have not yet completed terminal morphogenesis (Fig. S3). Comparing apical marker localization, SJ morphology, and the cytoplasmic and nuclear sizes of Su(H)-lacZ^+^ cells, we distinguished four stages of apical membrane morphogenesis.

In Stage 1 (left-most column in Fig. 3), enteroblasts are small, and their nuclei appear diploid; this stem-like appearance is consistent with the cells being in early stages of terminal differentiation. Stage 1 enteroblasts lack apical polarity, but they have formed SJ contacts with enterocytes. The few apical markers that are expressed, such as Moesin (Figs. 3A and C), do not show a polarized distribution; other apical markers, such as Karst and Mdu (Figs. 3B and 3C), are not detectable. The apex of Stage 1 enteroblasts contacts the basal terminus of enterocyte-enterocyte SJs, as revealed by localization of SJ components Tsp2A and Ssk (Figs. 3A and 3C). We interpret these SJ contacts to be nascent, point-like SJs, similar to those formed by the enteroblast in Figs. 2B and 2C.

In Stage 2 (Fig 3, second column from left), enteroblasts preferentially localize apical markers to their apex; this enrichment forms a bright plaque that is covered by broadened SJ contacts (Figs. 3A-C). We suggest that this apical plaque represents early stages of polarization since low levels of apical markers persist at other cortical regions. The enteroblast SJ grows to cover the cell’s entire apex; we infer that these Stage 2 SJs are similar to the SJ in Figs. 2D and 2E. With respect to morphology, some Stage 2 enteroblasts (e.g. Figs. 3B and 3C) are similar in cytoplasmic and nuclear size to Stage 1 enteroblasts; other Stage 2 enteroblasts are slightly larger, and their nuclei appear intermediate in ploidy between 2N stem cells and 32-64N, mature enterocytes (e.g. Fig. 3A).

In Stage 3 (Fig. 3, third column from left), the differentiating cells resemble immature enterocytes. Their cytoplasmic and nuclear volumes are larger than Stage 1-2 enteroblasts, yet smaller than mature enterocytes, and their low levels of β-galactosidase suggest that Su(H) enhancer activity was diminishing. We refer to these cells as pre-enterocytes. Apical markers—now highly expressed—localize to a conspicuous, concave structure that is covered by the broad, umbrella-shaped SJ sheet that we described above. These concave structures are the morphological hallmarks of Stage 3. With diameters up to 12 µm—roughly the diameter of a mature enterocyte—they often fill the apex of the pre-enterocytes. We were surprised to discover that they are topologically discontinuous with the midgut lumen (Video 3), another feature that is incompatible with radial intercalation. Instead, we conjecture that these structures are precursors of the pre-enterocytes’ future lumen-contacting surface. Hence, we designate them as Pre-Assembled Apical Compartments (PAACs).

In Stage 4, (Fig. 3, right column), pre-enterocytes finish integrating into the gut epithelium by acquiring a topology of cell-cell interfaces that is equivalent to mature enterocytes. Stage 4 cells are circumscribed (rather than covered as in previous stages) by SJs, and they now possess a lumen-contacting apical surface. The shapes of Stage 4 lumen-contacting surfaces and Stage 3 PAACs are highly similar, which suggests that the PAAC opens up to the gut lumen via remodeling of its overlying umbrella SJ into a ring. Stage 4 cells are smaller in cytoplasmic volume and nuclear size compared to mature enterocytes. In a final Stage 5 (Fig. S3D), the cell acquires its mature size and ploidy and everts its lumenal-apical surface to form a convex shape, thus completing terminal differentiation.

Altogether, this morphogenetic sequence reveals that midgut stem cell progeny do not form new SJs and an apical membrane via radial intercalation into the epithelial barrier. Rather, they form these barrier structures while still in the basal epithelium, protected by the mature barrier.

### Live imaging of enteroblast-enterocyte integration

We next asked whether live imaging corroborates the four-stage sequence (Fig. S3D) of enteroblast-enterocyte integration implied by fixed samples. To address this question, We performed continuous time-lapse imaging using Windowmount methodology, in which volumetric movies of physiologically functioning guts are captured in live animals through a window cut into the dorsal cuticle (Fig. 4A) (Martin et al., 2018). Prior analyses of stem cell clones in both fixed guts (He et al., 2019; de Navascués et al., 2012) and intravital live imaging (Koyama et al., 2020) suggest that most enteroblasts require >24 h to differentiate into enterocytes. Although this time frame is longer than the 16-20 h viability of animals during Windowmount (Martin et al., 2018), longitudinal imaging suggests that some cells differentiate in <24 h (Koyama et al., 2020). Therefore, we aspired to capture these faster cells.

**Figure 4:**
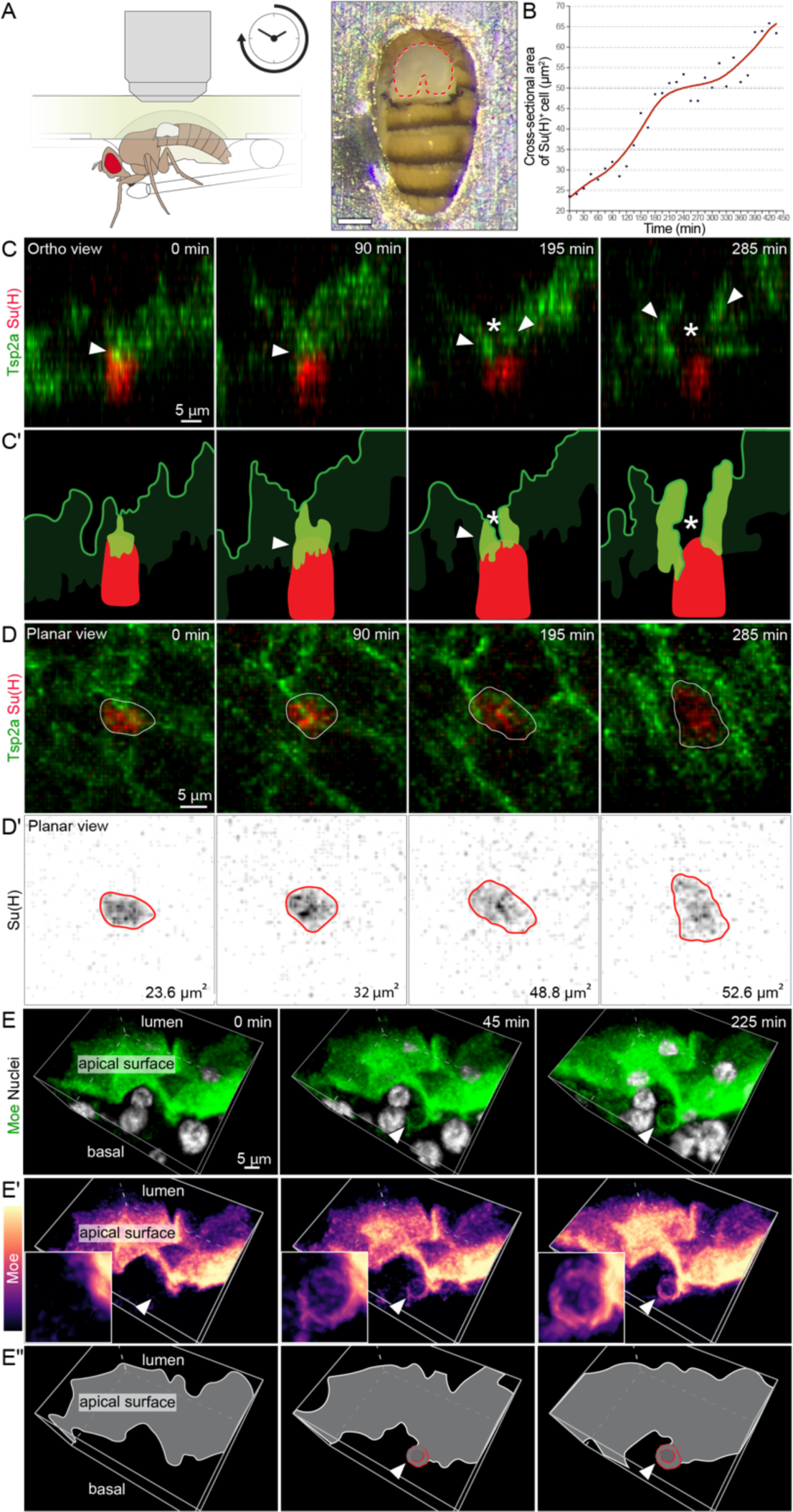
*In vivo* live imaging of SJ and PAAC dynamics supports the four-stage model of integration. (A) Continuous time-lapse imaging of midguts in live, feeding *Drosophila* was performed through a window in the dorsal cuticle. Adapted from Martin, 2018. (B) Cross-sectional area of the *Su(H)-mCherry*^+^ cell shown in Panels C and D and in Video 4. The increase in area implies that this cell is actively differentiating from enteroblast to enterocyte. (C-D) Live dynamics of SJ during enteroblast-enterocyte differentiation. Still frames are from a 7.25 h volumetric movie (Video 4) of a midgut expressing *GS5961-tsp2A::GFP* and *Su(H)-mCherry.* The cell analyzed in Panel B is shown in ortho view (C; apical at top) with corresponding line drawings (C’). In ortho view, arrowheads point to the SJ associated with this cell and asterisk denotes estimated location of putative PAAC. The planar view is shown in (D); panels are projections of serial confocal images. Numbers in the lower right corner of panels in D’ are cross-sectional areas of the cell at the given time points. SJ morphogenesis over time is visible in the ortho view: Between 0 min and 90 min, the SJ broadens over the cell’s apex. At 195 min, a hollow space (asterisk in C) develops along the SJ’s apical-basal axis. Between 195 and 225 min, the hollow space widens; both the hollow space and its surrounding SJ elongate along the lateral edges of the now-larger cell. (E) Live imaging of PAAC development. Still frames from a 3.75 h volumetric movie (Video 5) of midgut expressing the apical marker *moe::GFP* (green in E; magma LUT in E’) and the nuclear marker *ubi-his2av::RFP* (grayscale in E). Corresponding line drawings are in E’’. Arrowhead in E’ points to the area of PAAC formation, which is also shown as a close-up in the inset. At 0 min, the lumenal-apical surface appears as a lumpy blanket overlying the gut cell nuclei; no PAAC is visible. By 45 min, a putative PAAC has formed at the basal side of the lumenal-apical surface. By 225 min, the PAAC has become deeper and brighter. Insets in E’ show close-up views of the developing PAAC. Full genotypes for all panels in Table.

We first examined live dynamics of SJs. From nine movies of *GS5966>tsp2A::GFP; Su(H)-mCherry* midguts with durations from 7.25-20 h, we identified one movie in which an mCherry-expressing cell increased in cross-sectional area by nearly three-fold (Fig. 4B & 4D, and Video 4); this dramatic growth is an identifying feature of enteroblast-enterocyte differentiation. To determine whether the Tsp2A::GFP-labelled SJ associated with this cell exhibited dynamics consistent with the mechanism implied by Fig. 3, we analyzed an orthogonal view through the cell’s apical-basal axis (Fig. 4C). In the movie’s initial 105 min, the SJ that contacted the differentiating cell grew broader and ultimately covered the cell’s entire apex (Fig. 4C, arrow in 0- and 90-min panels; Video 4, 0-105 min). This broadening is consistent with the notion that SJs expand from discrete contact points in Stage 1 (Fig. 2C, Fig. 3A & 3B, and Video 1) to an umbrella shape in Stages 2 and 3 (Fig. 2E, Video 2, Fig. 3A & B, Fig. 5D, and Video 6). Between 120-285 min, a hollow space developed along the SJ’s apical-basal axis; simultaneously, the SJ extended along the lateral faces of the now-larger cell (Fig. 4C, arrows (SJ) and asterisks (hollowing) in 195- and 285-min panels; Video 4, 120-285 min). This hollowing and lateral extension are consistent with remodeling of the SJ from an overlying sheet in Stage 3 to a circumscribing ring in Stage 4 (Fig. 3A & 3B). Overall, these live SJ dynamics support the morphogenetic sequence suggested by the fixed analyses in Fig. 3.

**Figure 5.**
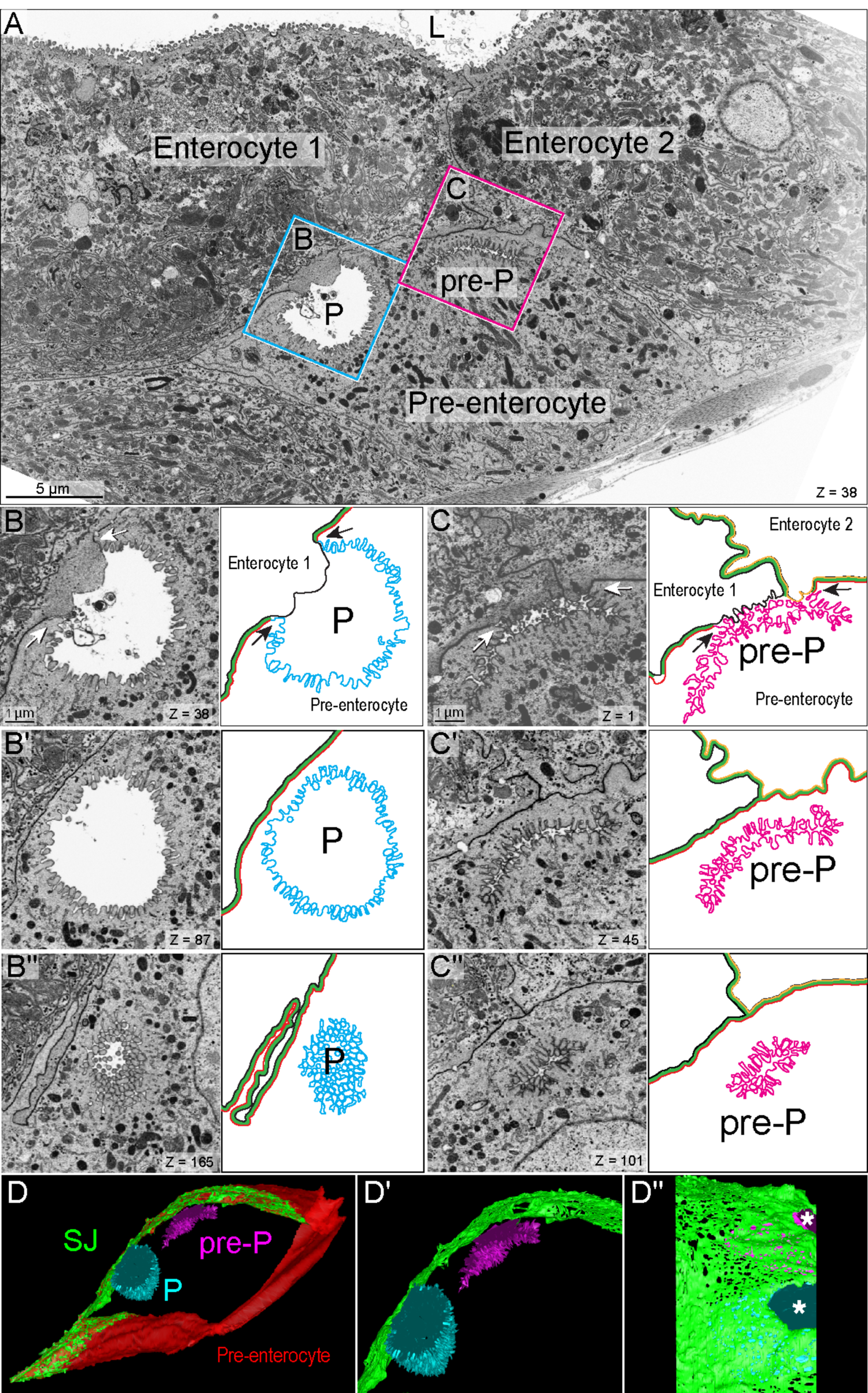
PAACs are intercellular lumens with split apical-basolateral polarity. (A) One slice of a representative, 200-slice FIB-SEM tomographic series. Series captures two mature enterocytes that contact the gut’s central lumen (L) and an underlying pre-enterocyte, which does not. An ellipsoid-shaped PAAC (P, cyan box) and an allantoid-shaped PAAC precursor (pre-P, magenta box) appear at the apex of the pre-enterocyte. See Figure S4. (B) Close-up of the PAAC in cyan box in A. Three FIB-SEM sections are shown next to cartoon representations. P indicates the PAAC lumen. In cartoons, the apical membrane of the pre-enterocyte is cyan, the basolateral membrane of the pre-enterocyte is red, the basolateral plasma membrane of enterocyte 1 is black, and the SJ between the pre-enterocyte and enterocyte 1 is green. In B, a gap in the SJ reveals that the PAAC is an intercellular lumen between the apical pre-enterocyte membrane and the basolateral enterocyte membrane (split polarity). Arrows point to the three-way boundary between the pre-enterocyte apical membrane, the enterocyte basolateral membrane, and the pre-enterocyte basolateral membrane. In B’ and B’’, the deep cytoplasmic invagination of the pre-enterocyte apical membrane forms most of the PAAC’s lumenal volume. (C) Close-up of the PAAC precursor in magenta box in A. pre-P indicates the PAAC precursor. In cartoons, the pre-enterocyte’s apical membrane is magenta, the pre-enterocyte’s basolateral membrane is red, the basolateral plasma membrane of enterocyte 1 is black, the basolateral membrane of enterocyte 2 is orange, and the SJ between the pre-enterocyte and enterocytes 1 and 2 is green. In C, a gap in the SJ reveals that the PAAC precursor is an intercellular lumen. Arrows point to two three-way boundaries between the pre-enterocyte apical membrane, the basolateral membrane of either enterocyte 1 or enterocyte 2, and the pre-enterocyte basolateral membrane. In C’ and C’’, the precursor’s slender, allantoid-shaped lumen arises through invagination of the pre-enterocyte’s convoluted apical membrane. In A-C, Z values in lower left of panels are slice numbers. (D) Volumetric rendering of 200 FIB-SEM sections, including the section in A. Apical membranes of the pre-enterocyte are cyan (PAAC) and magenta (PAAC precursor), basolateral membrane of the pre-enterocyte is red, and SJ between the pre-enterocyte and mature enterocytes is in green. Zoomed-in panels show the PAAC, PAAC precursor, and SJ in a cutaway view (D’) and a top-down view (D’’). In D and D’, the PAAC’s ellipsoid shape and the precursor’s allantoid shape are evident. In D’’, asterisks mark holes in the SJ resulting from separation of pre-enterocyte and enterocyte plasma membranes during PAAC formation. (Enterocyte membranes not shown.) See Video 6. Full genotype in Table 1.

We next examined live dynamics of the apical membrane using midguts that expressed the apical marker Moesin::GFP and the nuclear marker His2av::mRFP (Fig. 4E and Video 5). The Moesin::GFP-labelled lumenal-apical membranes appeared as a convoluted surface atop the gut cells’ nuclei because the apical surface of mature enterocyte is domed (*e.g.* Fig. 2A). Basal to the lumenal-apical membranes, we frequently observed GFP-labelled structures that were fainter in intensity and concave in shape. We conjecture that these structures are PAACs. While most PAACs did not dramatically change in shape or size during imaging, some PAACs became brighter and deeper over time (*e.g.,* Fig. 4E and Video 5). This evolution is consistent with Stages 3-4 of differentiation, during which PAACs initially form and subsequently become larger and more enriched for apical markers (Fig. 3, Stages 2-3). Overall, we conclude that live imaging of SJ and apical membrane dynamics provides additional support for the four-stage mechanism suggested by fixed tissues.

### PAACs are intercellular lumens formed by asymmetric pre-enterocyte apical membranes and enterocyte basolateral membranes

Our finding that PAACs are physically distinct from the gut’s lumenal-apical surface raises basic questions about the nature of these structures: Are they intracellular or intercellular? What is their relationship to the developing SJ? Does their apical polarity correspond to a mature brush border? To gain insight into these and other questions, we examined the PAACs’ ultrastructure in FIB-SEM tomographic series.

To identify putative PAACs, we first identified pre-enterocytes by looking for polyploid cells that lacked a visible lumenal-apical surface. We noticed that the apexes of such pre-enterocytes frequently contained membrane-bound, ellipsoid ultrastructures (Fig. 5A–cyan box, and Fig. 5B) whose shape and cellular position resembled PAACs. These structures enclosed prominent lumens that are distinct from the gut lumen, circumscribed by SJs, and lined with microvilli. The microvilli are densely arrayed, similar to brush border microvilli, but they are shorter, which suggests they are immature. Notably, Moesin, which outlines PAACs in confocal micrographs (Fig. 3 and Video 3), is a marker of microvilli in other Drosophila epithelia (Edwards et al., 1997; Lattner et al., 2019). We also found sausage-shaped (allantoid) ultrastructures that, like the ellipsoids, are lined with dense microvilli and circumscribed by SJs (Fig. 5A–magenta box, and Fig. 5C). The allantoids’ lumens are extremely slender, suggesting that they may be newly formed. Given these features, we conjecture that the ellipsoid ultrastructures are PAACs and the allantoid ultrastructures are their precursors.

We took advantage of our FIB-SEM series to investigate whether PAACs are intracellular compartments that develop within a pre-enterocyte, akin to the large apical endosomes (vacuolar apical compartments/apicosomes) observed in mammalian cells (Gilbert and Rodriguez-Boulan, 1991; Taniguchi et al., 2017; Vega-Salas, 1988) or intercellular compartments that develop between a pre-enterocyte and its mature enterocyte neighbors. We selected series that captured the near-complete volume of individual PAACs or PAAC precursors, and we analyzed their membrane topologies slice-by-slice (Figs. 5B and 5C). We also generated a volumetric rendering of a 200-slice FIB-SEM tomographic series that contained a PAAC, a PAAC precursor, and their associated pre-enterocyte within a tissue volume of 40.2 µm x 23.9 µm x 8 µm (Fig. 5D and Video 6).

These analyses invariably uncovered a region in which pre-enterocyte and mature enterocyte membranes separate from each other to form the PAAC’s lumenal space (Fig. 5B and Video 6). Thus, PAACs are intercellular. The PAAC lumen is surrounded by an expansive SJ that adheres the pre-enterocyte to mature enterocytes; this SJ separates the PAAC from the gut’s central lumen. The ultrastructure of the PAAC-associated SJ is consistent with the SJs we observed in immunostained, Stage 3 pre-enterocytes (Fig. 3 and Video 3). PAAC precursors—even very small ones—also comprise an intercellular lumen surrounded by an expansive SJ. We did not observe any microvilli-lined compartments that were entirely intracellular. The similar topology of PAACs and PAAC precursors suggests that PAACs initiate via de-adhesion of apposing plasma membranes rather than fusion of an intracellular compartment with the SJ.

### PAACs’ split apical/basolateral polarity is unique for a lumen-encompassing structure

PAAC-forming pre- and mature enterocytes make markedly unequal contributions to the PAAC’s overall morphology. The pre-enterocyte plasma membrane represents the vast majority of a PAAC’s total surface area. It folds inward at nearly 180 degrees (arrow in Fig. 5B) to create a deep invagination into the pre-enterocyte cytoplasm. This invagination accounts for most of the PAAC’s lumenal volume and evokes a scenario in which PAAC development is driven by inward folding of the pre-enterocyte plasma membrane. In contrast to the structured folds of the pre-enterocyte membrane, the mature enterocyte membranes are amorphous and rest like a blobby lid atop the PAAC lumen (Fig. 5B). These differences are even more extreme in PAAC precursors (Fig. 5C).

The contrast between the structured pre-enterocyte membrane and the amorphous enterocyte membrane corresponds to a second—unprecedented—asymmetry of PAACs: their lumenal polarity is split. The PAAC’s pre-enterocyte membrane is apical while the partner enterocyte membrane(s), which lacks apical markers (Video 3) and microvilli (Figs. 5B & 5C, and Video 6), is by default basolateral. To our knowledge, this combination has not previously been reported for any epithelial lumen either in cell culture or *in vivo*. Rather, epithelial lumens to date have been either uniformly apical (Blasky et al., 2015; Datta et al., 2011; O’Brien et al., 2002), or, in some rare, experimentally induced cases, uniformly basolateral (Lowery et al., 2009; Wang et al., 1990). However, these prior studies examined lumens that form between cells at similar differentiation states; we conjecture that the PAACs’ unique asymmetries derive from their origin between cells at distinct differentiation states.

### Enteroblasts must form SJs to integrate and mature into enterocytes

Our data reveal that as stem cell daughters differentiate, they initiate epithelial integration by forming new, sheet-like SJs with mature neighbor cells. What happens when SJ formation is blocked? To examine this question, we generated “SJ-less” enteroblasts and assessed their ability to integrate and differentiate. We inhibited expression of the SJ component *ssk* specifically in enteroblasts by using the enteroblast driver *Su(H)-GAL4* to express a *UAS-sskRNAi* transgene under control of temperature-sensitive *GAL80^ts^* (McGuire et al., 2003) (genotype henceforth referred to as *Su(H)^ts^>sskRNAi*). A *UAS-GFP* transgene was also included to identify the RNAi-expressing cells. The *sskRNAi* hairpin was expressed from days 0-4 of adult life, after which midguts were harvested and analyzed. To confirm that *sskRNAi* prevented SJ formation, we performed immunostaining for another SJ component, Coracle. Whereas Coracle localized to the apex of *Su(H)^ts^*-expressing control cells, it localized to the cytoplasm of *Su(H)^ts^>sskRNAi* cells (Figs. 6A & 6B). This redistribution implies that *sskRNAi* expression prevents proper formation of SJs.

**Figure 6.**
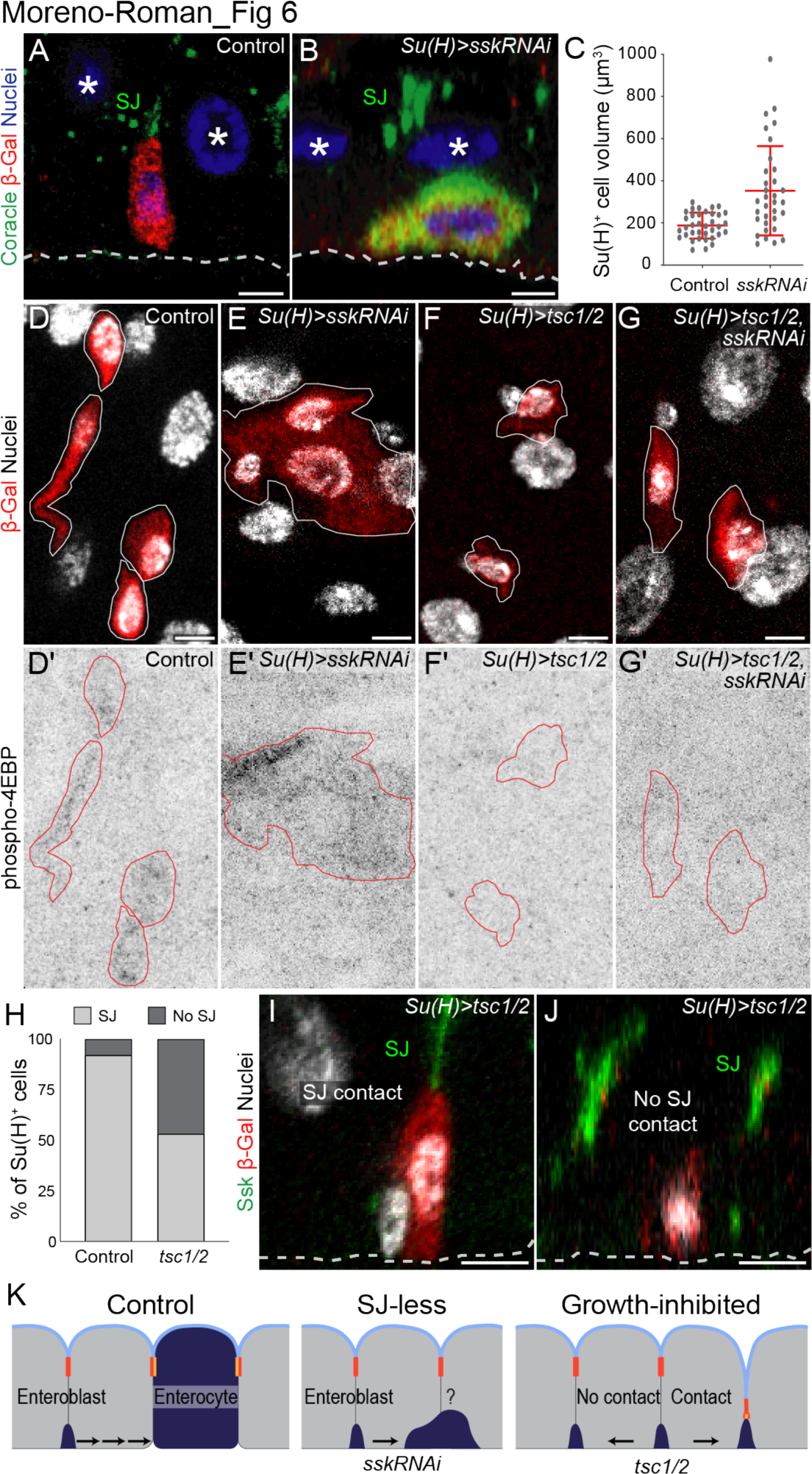
Cells must form SJs and grow in order to integrate. (A-C) Blocking SJ formation prevents integration but not growth. (A) Cross-sectional view of control *Su(H)*^+^ cell (red, *Su(H)-lacZ*). The cell’s apex has formed contacts with the basal tip of the SJ (green, Coracle) between neighbor enterocytes. (B) Cross-sectional view of *Su(H)^ts^>sskRNAi*. The cell expresses *Su(H)-lacZ* (red) and high levels of Coracle (green), which localizes to the cytoplasm. It does not contact either enterocyte-enterocyte SJs or the gut lumen. (C) Measurements of cytoplasmic volumes show that *Su(H)^ts^>sskRNAi* cells become larger in size compared to control *Su(H)* cells. (D-G) Growth of *Su(H)^ts^>sskRNAi* cells requires Tor pathway activation. Planar views of all midgut cell nuclei (grayscale, DAPI) and *Su(H)-lacZ* (red) are shown above corresponding images of phospho-4EBP immunostain (D’-G’, inverted grayscale). *Su(H)-lacZ*-labelled cells are outlined in white (D-G) and red (D’-G’). Control *Su(H)* cells (D) and *Su(H)^ts^>sskRNAi* (E) cells are phospho-4EBP^+^ (D’, E’). Tor-inhibited, *Su(H)^ts^>tsc1/2* (F) and *Su(H)^ts^>sskRNAi, tsc1/2* (G) are not (F’, G’). (H-J) Growth-inhibited enteroblasts arrest at initial stages of integration. The frequency with which *Su(H)^ts^>tsc1/2* cells (*Su(H)-lacZ*, red) contact the gut SJ network (Ssk, green) is reduced to 53% from the control frequency of 92% (H). Both *Su(H)^ts^>tsc1/2* cells that contact SJs (I) and those that do not (J) fail to reach the gut’s lumenal surface. N=3 control midguts (214 enteroblasts) and 3 *Su(H)>tsc1/2* midguts (146 enteroblasts). All scale bars, 5 µm. Full genotypes in Table 1. (K) Cartoon summary: (Left) Control. Integration requires both cell growth and SJ formation. (Middle) Blocking SJ formation prevents integration but does not halt growth. (Right) Growth inhibition arrests cells in Stages 0-1 of integration.

*Su(H)^ts^>sskRNAi* cells categorically failed to integrate into the gut epithelium. Instead of reaching the gut’s central lumen, these cells accumulated beneath the gut’s SJ network (Figs. 6A & 6B). Although this basal localization is typical of enteroblasts, *Su(H)^ts^>sskRNAi* cells did not arrest in an enteroblast state. Rather, they grew in volume and endoreplicated their nuclei (Fig. 6C and Figs. S5A & S5B), two behaviors characteristic of pre-enterocytes (Fig. 3, Stages 3 & 4). Yet unlike pre-enterocytes, *Su(H)^ts^>sskRNAi* cells did not adopt a cuboidal or columnar shape and instead became blob-shaped. Similar behaviors were exhibited by MARCM-generated stem cell clones (Lee and Luo, 1999) that were genetically null for SJ components *tsp2A* (Chen et al., 2018) and *mesh* (Fig. S5E & S5F). Thus even as SJ-less cells achieve mature size, they do not become part of the epithelium.

The indeterminate morphology of *Su(H)^ts^>sskRNAi* cells was accompanied by inappropriate, mixed expression of fate-specific transcription factors. Whereas the stem cell/enteroblast transcription factor Sox100B (Doupé et al., 2018; Jin et al., 2020; Meng et al., 2020) was expressed by both control *Su(H)* and *Su(H)^ts^>sskRNAi* cells (Figs. S5A & S5B), the enterocyte transcription factor Pdm1 (Dantoft et al., 2013; Korzelius et al., 2014; Lee et al., 2009) was absent from control *Su(H)* cells yet expressed in *Su(H)^ts^>sskRNAi* cells (Figs. S5C & S5D). Similar, mixed patterns of marker expression were observed by Xu and colleagues upon enteroblast-specific depletion of the SJ component *tsp2A* (Xu et al., 2019).

Altogether, these findings imply that SJ-less cells become trapped in an abnormal, hybrid cellular state in which the distinct features of enteroblasts and enterocytes co-exist abnormally.

### Cell growth is required for integration, independent of SJs

Having found that growth is not sufficient for cells to integrate, we asked whether integration requires cell growth. Cell growth during the enteroblast-enterocyte transition is controlled by the Tsc/Rheb/Tor pathway (Amcheslavsky et al., 2011; Kapuria et al., 2012; Nie et al., 2015; Quan et al., 2013; Xiang et al., 2017). Tor pathway activation in enteroblasts can be visualized by immunostaining for the phosphorylated isoform of the Tor kinase substrate eIF4E Binding Protein (phospho-4EBP) (Kapuria et al., 2012). When Tor is inactivated via overexpression of its inhibitor, Tsc1/2, 4EBP is not phosphorylated (Fig. 6F) and differentiation-associated growth is blocked (Kapuria et al., 2012).

We found that growth of SJ-less cells, like growth of normal enteroblasts, depends on Tor. Phospho-4EBP immunostaining showed that *Su(H)^ts^>sskRNAi* cells are Tor-activated, akin to *Su(H)* control cells (Figs. 6D & 6E). When we conditionally overexpressed *tsc1/2* in either control cells (*Su(H)^ts^>tsc1/2*) or *ssk* knockdown cells (*Su(H)^ts^>sskRNAi, tsc1/2*), we abrogated phospho-4EBP and inhibited cell growth (Figs. 6F & 6G).

We next examined whether Tor inactivation and consequent growth inhibition affects the ability of cells to integrate. We assessed SJ formation in *Su(H)^ts^>tsc1/2* cells by immunostaining guts for Ssk and the *Su(H)-lacZ* reporter and determining whether the Ssk signal contacted the apex of β-galactosidase-labelled cells. Whereas 92% of control *Su(H)* cells formed SJs, only 53% of *Su(H)^ts^>tsc1/2* cells did (Fig. 6H). Revealingly, no *Su(H)^ts^>tsc1/2* cells progressed beyond Stage 1 (Fig. 6I). Thus, SJ initiation is not sufficient for integration to progress; enteroblast growth is also necessary. While the precise contribution of growth is currently unclear, it may fuel expansion of the umbrella SJ or to initiate PAAC formation.

### Organ-scale impacts of blocked cellular integration

Organ renewal requires that new cells integrate successfully into the epithelium. When cell integration is blocked, what are consequences to organ-scale cellular equilibrium? We first asked whether blocking integration causes undifferentiated cells to accumulate abnormally in the tissue (Fig. 7A). When animals are maintained under stable, *ad libitum* conditions, Su(H)^+^ cells typically comprise ∼10% of total cells in the midgut R4 region (Bonfini et al., 2021; O’Brien et al., 2011; Viitanen et al., 2021). This proportion essentially doubled when we blocked integration by inhibiting new SJ formation (19.1% ± 5.5% of total cells in *Su(H)^ts^>sskRNAi* guts; 9.3% ± 2.8% in control guts).

**Figure 7.**
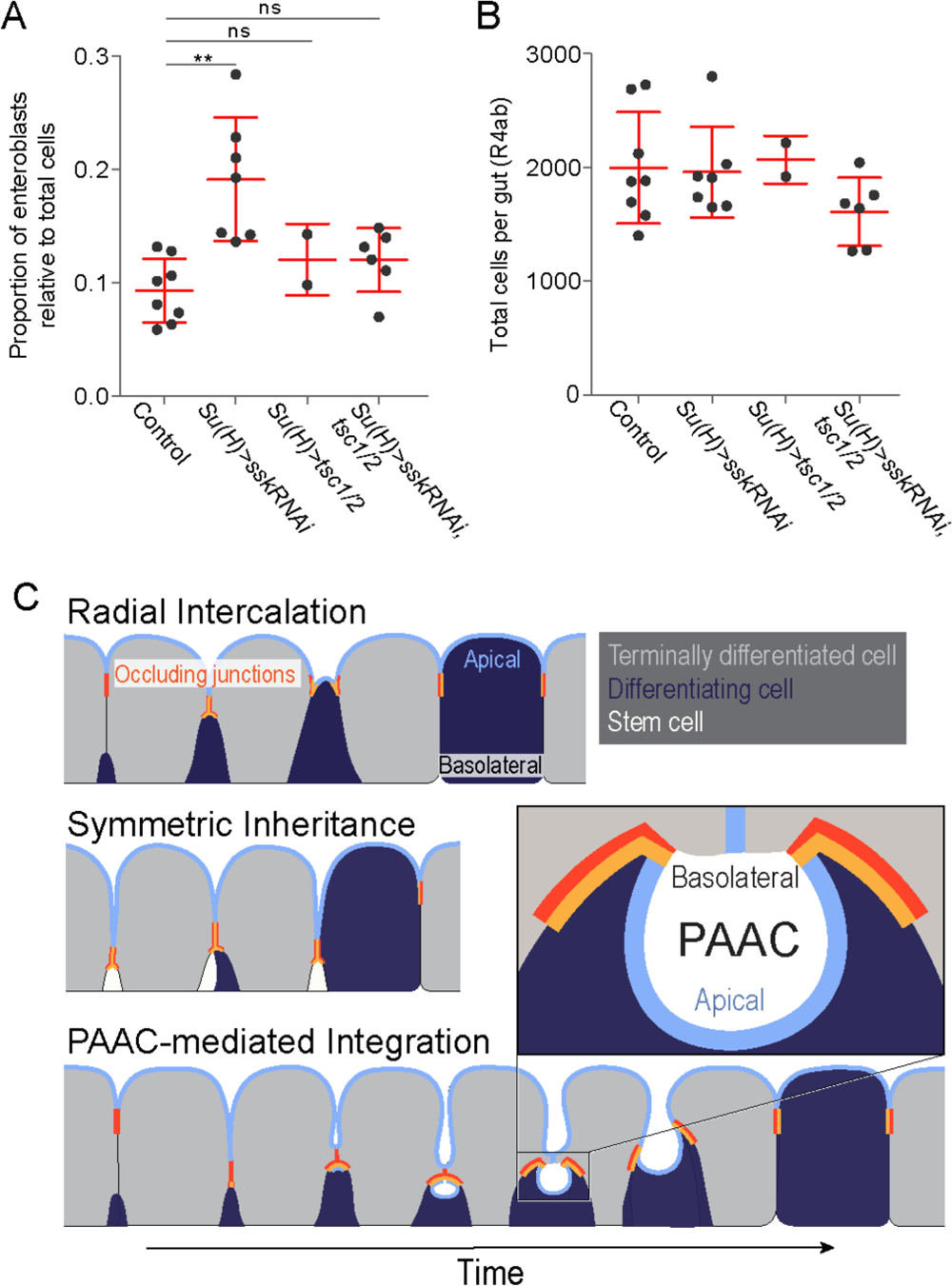
Impact of blocked midgut cell integration on organ-scale cell equilibrium. (A) Proportion of Su(H)^+^ cells in midguts with blocked cell integration. Plots show the percentage of *Su(H)-lacZ*^+^ cells relative to total cells in the R4ab region of midguts with the indicated genotypes. Cell integration was blocked by inhibiting either new SJ formation (*Su(H)^ts^>sskRNAi*), cell growth (*Su(H)^ts^>tsc1/2*), or both (*Su(H)^ts^>sskRNAi, tsc1/2*) between days 4-8 of adult life. Each data point represents one midgut. Red lines show Means ± S.D for each condition: *Su(H)* control –0.09 ± 0.03 cells; *Su(H)^ts^>sskRNAi* – 0.19 ± 0.05; *Su(H)^ts^>tsc1/2* – 0.12 ± 0.032 cells; *Su(H)^ts^>sskRNAi, tsc1/2* – 0.12 ± 0.028 cells. (B) Total numbers of midgut cells remain constant when cell integration is blocked. Plots show total counts of DAPI-labeled nuclei in the R4ab regions of midguts analyzed in Panel A. Each data point represents one midgut. Red lines show Means +/- S.D for each condition: *Su(H)* control – 1997 ± 489 cells; *Su(H)^ts^>sskRNAi* – 1960 ± 398; *Su(H)^ts^>tsc1/2* – 2067 ± 210 cells; *Su(H)^ts^>sskRNAi, tsc1/2* – 1611 ± 299 cells. (C) Three mechanisms to incorporate stem cell progeny into a mature epithelium. Only PAAC-mediated integration enables the differentiating cell to form new barrier structures (apical plasma membrane and occluding junctions) while still sheltered by the mature occluding junction barrier.

By comparison, the proportion of Su(H)^+^ cells remained nearly normal when we blocked integration by inhibiting cell growth (12.0 ± 2.8% in *Su(H)^ts^>tsc1/2* guts). Concomitant inhibition of both SJ formation and cell growth resembled growth inhibition alone (12.0 ± 3.2% of total cells in *Su(H)^ts^>sskRNAi, tsc1/2*guts were Su(H)-lacZ^+^). Thus, whether integration-blocked cells accumulate in the tissue depends on the means through which integration was blocked. One possible reason may be differences in differentiation state of the integration-blocked cells. Early-stage enteroblasts, which still adhere strongly to their mother stem cell, can repress subsequent mother cell divisions (Choi et al., 2011), and Tsc1/2 overexpression—but not SJ inhibition—arrests differentiation at an early stage ((Kapuria et al., 2012) and Figs. 6C & S5C-G). This early-stage arrest may enable growth-inhibited cells to repress production of additional daughter cells.

We next examined whether blocking integration alters the organ’s total number of cells (Fig 7B). Comprehensive counts of DAPI-labeled nuclei in the midgut R4ab region revealed that total cell number remains normal when new cells cannot integrate, even for guts in which integration-blocked cells accumulate abnormally (1997 ± 489 cells in control *Su(H)^ts^* guts compared 1960 ± 398, 2067 ± 210 cells, and 1611 ± 299 cells in *Su(H)^ts^>sskRNAi*, *Su(H)^ts^>tsc1/2*, and *Su(H)^ts^>sskRNAi, tsc1/2* guts, respectively). We speculate that feedback mechanisms inherent to organ-scale control of total cell number (Akagi et al., 2018; Jin et al., 2017; Liang et al., 2017) ‘sense’ unintegrated cells and exert a compensatory effect on cellular equilibrium.

## DISCUSSION

Epithelial organs maintain a leakproof barrier between the interior body and the external environment even while continuously replacing the cells that directly contact this environment. In many barrier epithelia, these replacement cells derive from basal stem cells and are born without a lumenal-apical surface or occluding junctions, two structures that are essential for barrier integrity. Consequently, daughters must generate these structures *de novo* and integrate into the barrier as they differentiate.

We examined this process at ultra-fine spatial resolution during physiological turnover of the *Drosophila* intestinal epithelium. Our analyses led to a previously undescribed mechanism that we term PAAC-mediated integration (Fig. 7C): The new cell forms a broad, umbrella-shaped SJ that serves as a transient niche for biogenesis of the cell’s future lumenal-apical surface (the PAAC).

When the new cell is sufficiently mature, the umbrella SJ retracts and the PAAC lumen fuses with the gut lumen, exposing the cell’s apical membrane to the external environment. In contrast to a prior model of radial intercalation (Fig. 7C), PAAC-mediated integration enables stem cell daughters to form barrier structures in a space sheltered from the contents of the gut lumen—a potentially crucial safeguard for an epithelium that is simultaneously physiologically active and continuously renewing.

### PAAC architecture: Implications for epithelial lumen formation

Lumens are defining features of epithelial tubes, and the molecular and cellular events that drive lumen formation are a topic of intense interest. Our current understanding of lumen formation comes from studying epithelial cells that are at similar states of differentiation (Blasky et al., 2015; Datta et al., 2011; Overeem et al., 2015; Sigurbjörnsdóttir et al., 2014). PAACs provide a first, fine-grained example of how lumens form between cells that are at disparate states of differentiation. This fate difference likely underlies the PAACs’ two distinctive characteristics, structural asymmetry and split polarity. Below, we speculate how these PAAC-specific characteristics may shed new light on lumen-forming mechanisms in general.

PAACs are, to the extent we can determine, the first type of intercellular lumen that exhibits split polarity––the membranes that form PAACs alternate between apical identity (the pre-enterocyte) and basolateral identity (mature enterocytes) (Figs. 3, 5 & 7C; and Video 6). By comparison, all epithelial lumens of which we are aware are normally enclosed by membranes that are exclusively apical (Blasky et al., 2015; Datta et al., 2011; Overeem et al., 2015; Sigurbjörnsdóttir et al., 2014). Our finding that PAACs’ mature enterocyte membranes do not form a secondary apical domain is surprising because cells that contact multiple lumens in developing epithelia form a corresponding apical domain for each lumen (Alvers et al., 2014; Bagnat et al., 2007; Bryant et al., 2010). One possible explanation is that terminally differentiated epithelial cells actively repress secondary apical domains whereas epithelial cells in developmental contexts do not. Another, non-exclusive, possibility is that all lumens transiently exhibit split apical/basal polarity at their earliest stage—one lumen-forming cell initiates an apical domain prior to the others—but that this stage is, in most cases, extremely short-lived, so it has not been detected previously.

PAACs’ second striking feature is their extreme structural asymmetry: pre- and mature enterocyte membranes, despite being bonded at their edges by the same SJ, acquire shapes that are extreme opposites. The pre-enterocyte PAAC membrane, which grows dramatically and invaginates deeply into the cytoplasm of the differentiating cell, convolutes into sharp folds and broad curves. This structure, which is superimposed onto the membrane’s fine-scale microvillar folds, sets the volume of the PAAC. The mature enterocyte membranes, by contrast, appear largely passive. They sit like a lid atop the neck of the pre-enterocyte invagination and do not appear to morph or expand during differentiation (*e.g.,* compare the nascent and advanced PAACs in Fig. 5 and Video 6).

These two features, split apical/basolateral polarity and structural asymmetry, provide insight into opposing models of lumen formation. The structural asymmetry of PAACs is incompatible with the prevailing model of lumen formation, in which hydrostatic pressure drives lumen growth (Chan et al., 2019; Dasgupta et al., 2018; Dumortier et al., 2019; Ruiz-Herrero et al., 2017; Yang et al., 2021), because lumens generated by hydrostatic pressure are uniformly convex (Vasquez et al., 2021). Rather, this asymmetry evokes a recently proposed alternative mechanism in which expansion of apical membrane surface drives lumen growth in a pressure-independent manner (Vasquez et al., 2021). Indeed, since growth of the PAAC lumen is accounted for by apical surface expansion of a single cell, PAACs may provide an informative case study of apical surface-driven growth. Identifying the molecular signals that target and stabilize new PAACs will aid in exploring these scenarios.

### A trade-off between barrier integrity and junction-forming efficiency

In both PAAC-mediated integration and radial intercalation, single cells assimilate into an epithelium through basal-to-apical movement. Why do distinct mechanisms exist to reach the same cellular endpoint? We speculate that, in general, basal-to-apical assimilation requires a trade-off between integration speed and barrier integrity. A given mechanism may favor one of these qualities at the expense of the other. In principle, a spectrum of mechanisms enables tissues to employ the mechanism that is best suited to their specific biological context.

In this schema, radial intercalation is rapid and parsimonious. It occurs over time scales of minutes or a few hours. New junctions initiate within the pre-existing junctional network and expand directly into their final morphology, an apico-lateral band. Intriguingly, all examples of radial intercalation described in the literature take place in developing epithelia (Merzdorf et al., 1998; Deblandre et al., 1999; Stubbs et al., 2006; Voiculescu et al., 2007; McMahon et al., 2008; Campbell et al., 2010). Because embryos themselves are housed in a protective environment (such as an egg or a womb), cells in embryonic tissues can display immature junctions and incipient microvilli at the organ’s apical surface without risking exposure to the external environment.

PAAC-mediated integration is slower and, since it involves building and then retracting a temporary scaffold, likely less efficient. We estimate that typical time frames for PAAC-mediated integration are >24 h, based on changes in nuclear size of stem cell daughters over time (Koyama et al., 2020). (Indeed, this >24 h time frame presented a challenge for Windowmount live imaging, which typically yields movies ∼8-20 h in duration.) A differentiating cell may require this time to construct the umbrella SJ and microvilli-lined PAAC—large structures that will undergo extensive remodeling in subsequent stages. The indirect, more complex nature of PAAC-mediated integration provides an additional layer of protection for differentiating cells—a potentially worthwhile tradeoff for a functionally active organ that continuously processes substances from the external environment.

How do SJs guide differentiating cells into the epithelial sheet? One appealing notion is that they exert myosin-based pulling forces that draw differentiating cells toward the lumen (Varadarajan et al., 2019; Yu and Zallen, 2020). We were, however, unable to identify any impact on integration following enteroblast-specific inhibition of Rho kinase (Rok; data not shown). Since Rok is an essential activator of myosin contractility, this finding implies that myosin-based forces are not required in the integrating cell. A second possibility is that myosin-based pulling forces, or some other cytoskeletal regulator, is required in mature neighbor cells to aid the basal-to-apical movement of differentiating cells. Since SJs bond together mature and differentiating cells, they are well-positioned to coordinate these two cell types during integration. Finally, SJs may serve to polarize growth and/or cytoskeletal assembly along the apical-basal axis of the differentiating cell (Madara, 1987).

### Diverse epithelial architectures may use a diversity of cell assimilation mechanisms

Numerous barrier epithelia, including mammalian trachea, cornea, and olfactory lining, have a cellular organization similar to the fly gut (Chepko and Dickson, 2003; Chepko and Smith, 1997; Cotsarelis et al., 1989; Evans and Moller, 1991; Leung et al., 2007; Michael J. Evans, 2001; Rock et al., 2009; Sekiya et al., 1988; Tsujimura et al., 2002). All these tissues are renewed by basally localized stem cells that lack occluding junctions and lumenal-apical surfaces. Their daughter cells thus all face the same architectural challenge of integrating seamlessly into the barrier while they differentiate. Whether they overcome this challenge through PAAC-mediated integration, like the fly gut, or through some other, perhaps as-yet-undefined, mechanism will be an interesting question for future investigation.

In considering how epithelial architecture affects new cell integration, it is notable that two of the best-understood barrier epithelia, mammalian intestine and lung alveoli, sidestep the challenge of barrier integration entirely. In these tissues, stem cells possess both occluding junctions and lumenal-apical surfaces, and daughter cells symmetrically inherit these structures from their mother (Fig. 7C) (DeMaio et al., 2009; Fleming et al., 2007; Jinguji and Ishikawa, 1992; McKinley et al., 2018). Symmetric inheritance is morphogenetically parsimonious, but it requires the abscission of existing junctional septa and creation of new septa at the new daughter-daughter interface. Since this remodeling happens at the lumenal surface, it might produce potential weak points in the barrier. Thus, at first glance, the fact that mammalian intestine and lung use symmetric inheritance to assimilate new cells seems at odds with the idea that physiologically active epithelia need extra safe-guards to protect barrier integrity.

We speculate, however, that this potential risk is mitigated by these tissues’ particular architecture; namely, deeply recessed niches—intestinal crypts and terminal alveolar endbuds—in which the lumen-exposed stem cells reside. Crypts and endbuds are secluded from bulk lumenal flow, which provides their resident stem cells with built-in protection that stem cells in other epithelia lack. This built-in protection conceivably affords stem cell daughters the simplicity of directly inheriting barrier structures from their mother cell. In this light, PAACs may be seen as a cellular-scale solution for epithelia that lack recessed stem cell niches, the tissue-scale solution for protecting new cells. As such, our findings spotlight the intimate relationship between physiological function, organ form, and cellular differentiation and morphogenesis.

## MATERIALS and METHODS

### Drosophila husbandry

Mated adult female flies were used in all experiments. Crosses utilizing the TARGET system (GAL4/GAL80^ts^) were performed at 18°C (McGuire et al., 2003). Upon eclosion, adult animals were temperature shifted to 29°C for 4 days to inactivate GAL80^ts^ and induce GAL4-mediated expression. Midguts were harvested for immunostaining 4 days after induction. Flies used for immunofluorescence were raised on standard molasses medium at 18°C. Upon eclosion, they were shifted to 29°C. Midguts were harvested for immunostaining 4 days after eclosion.

### Immunohistochemistry and sample preparation for confocal microscopy

Dissected guts were fixed in 4% formaldehyde in PBS (pH 7.4) at room temperature for 1 hour, immunostained, and mounted as previously described (O’Brien et al., 2011). Primary antibodies: mouse anti-Armadillo (1:100, DSHB N2 7A1), rabbit anti-Snakeskin (1:1000, gift from Furuse lab), mouse anti-Coracle (1:50, DSHB C615.16), mouse anti-β-galactosidase (1:400, Promega Z3781), rabbit anti-Phospho4EBP1 (1:500, Cell Signaling). Secondary antibodies: donkey anti-rabbit IgG conjugated to Alexa 555 and donkey anti-mouse IgG conjugated to Alexa 647 (1:1000, Invitrogen A-31572 and A-31571, respectively). Nuclei were stained with DAPI (LifeTechnologies D1306).

Samples were incubated with primary antibody overnight at 4°C in PBT (PBS with 3% Triton X-100 (Sigma-Aldrich X100-100 mL)) with 5% NGS (Capralogics GS0250), washed 3 times in PBT, then incubated with secondary antibody for 4 hours at room temperature in PBT with 5% NGS. Samples were mounted in ProLong (LifeTechnologies P36984) and stored at −20°C until imaging.

### Induction of MARCM clones

Heat-shock MARCM clones (Lee and Luo, 1999) were generated by collecting adult flies 12-24 hours post-eclosion and performing two 45-min, 37°C heat shocks separated by a 8-min chill on ice. Flies were returned to 25°C for 4 days, then dissected and analysed.

### Fixed sample imaging

Fixed samples were imaged on a Leica SP8 WLL confocal microscope with a 63x HC PL APO CS2 oil objective. Serial optical sections were taken at 0.5 µm intervals through the entirety of whole-mounted, immunostained midguts.

### Quantitation of SJ-contacting and non-contacting cells

Enteroblasts in the midgut R4ab region (also known as P1/2) (Buchon et al., 2013; Marianes and Spradling, 2013; O’Brien, 2013) were visualized and counted using ImageJ/Fiji (Schindelin et al., 2012). The R4ab region was identified using morphological landmarks. SJs were identified by immunostaining for the SJ component Snakeskin. Su(H)-lacZ^+^ (β-Gal^+^) cells were recognized as enteroblasts by visual inspection. To categorize enteroblasts as SJ-contacting or non-contacting, each enteroblast was analyzed through Fiji Orthogonal View. Enteroblasts were defined as SJ-contacting cells if the β-Gal^+^ signal was juxtaposed and/or displayed overlap with apical Snakeskin signal in XY, XZ and YZ planes. Enteroblasts were defined as SJ non-contacting cells if they lacked these criteria.

### Measurements of cell volume

To measure volumes of *Su(H)-lacZ^+^* cells, tissues were fixed, immunostained using anti-β-galactosidase antibody, mounted, and subjected to volumetric confocal imaging. After initial processing in Fiji, files were imported to Bitplane Imaris v.8.7. Volumes of *Su(H)-lacZ^+^* cells were determined by creating a surface for each cell using the Imaris contour tool and then measuring the enclosed volume.

### Ovary dissection and staining

Egg chambers were dissected in phosphate-buffered saline (PBS pH 7.4) + 0.1% Triton X-100 and incubated for 2 hours in 1mM latrunculin B (LatB; Sigma). They were then fixed 20 min in 4% PFA (in PBS pH 7.4), incubated 2 hours in a 1:250 dilution of TRITC-conjugated phalloidin (Molecular Probes, Eugene, OR), and subsequently imaged on a Zeiss LSM 700 confocal microscope.

### Total cell and total enteroblast counts

To perform total cell counts and total enteroblast counts of R4ab, confocal image stacks were digitally isolated in Fiji. Bitplane Imaris was used generate three-dimensional organ reconstructions, and individual cells were comprehensively counted by mapping signals for DAPI (for total cell counts) or Su(H)-lacZ (for total enteroblast counts) to Imaris surface objects. Imaris-recognized surfaces were confirmed through visual inspection and manually adjusted when needed for accuracy.

### *In vivo* live imaging and movie analyses

Live imaging was performed on 2-3 day old adult females as described previously (Martin et al., 2018), with the following modifications: For Video 4, 20 nM RU486 and 10 μg/mL Isradipine were added to the imaging media to induce GeneSwitch5966 expression and reduce intestinal peristalsis, respectively. For Video 5, 10 μg/mL Isradipine was added to the imaging media to reduce intestinal peristalsis.

Videos were acquired with a LSM Leica SP5 with a HCX APO L 20x/ 1.00W lens, controlled by LAS AF software. Confocal sections were taken every 15 mins with z-steps of 1.01μm (Video 4) and 2.98 μm (Video 5). Videos were processed on a Windows computer (Windows 10 Education) with a 3.70 GHz quad-core Intel Xeon processor and 256 GB memory. Videos were initially processed in Fiji and subsequently visualized in volumetric format and analyzed in Bitplane Imaris. For Video 4, the following Fiji plugins were applied: 1) Stack Sorter (https://www.optinav.info/Stack-Sorter.htm), to correct the alignment of out-of-order slices captured during a peristaltic contraction, 2) StackReg (Arganda-Carreras et al., 2006) to correct for whole-organ X-Y movements, 3) Correct 3D Drift (Parslow et al., 2014) to correct for global volume movements, and 4) TrakEM2 (Cardona et al., 2012) to perform manual X-Y alignment for slices that could not be registered automatically. The latter 3 plugins were applied iteratively as needed. For Video 5, Stack Sorter, StackRed, and Correct 3D Drift were used. The latter two plugins were applied iteratively as needed.

### Measurements of cross-sectional cell area in live movies

The cross-sectional area of the integrating Su(H)-mCherry^+^ cell at each timepoint in Video 4 was determined as follows: The cell’s largest cross-sectional plane at each time point was identified by visual inspection. FIJI Measure was used to manually outline the mCherry signal in this plane and to measure the enclosed area.

### Sample preparation for FIB-SEM

Fly guts were dissected in PBS and immediately processed as previously described (Daniel et al., 2018; Kolotuev, 2014). Briefly, the samples were fixed in 1% formaldehyde, 2.5% glutaraldehyde in 0.1M phosphate buffer (PB) for 2 hours at room temperature, then incubated for 1 hour in 2% (wt/vol) osmium tetroxide and 1.5% (wt/vol) K4[Fe(CN)6] in PB followed by 1 hour in 1% (wt/vol) tannic acid in 100 mM cacodylate buffer, then 30 minutes in 2% (wt/vol) osmium tetroxide in water followed by 1% (wt/vol) uranyl acetate for 2 hours at room temperature. After the dehydration cycles, samples were embedded in Epon-Araldite mix. Samples were flat embedded to assure the targeting of the region of Interest during the sectioning step.

### Sample preparation for CLEM

To preserve native fluorescence for correlative light/electron microscopy, samples were subjected to high-pressure freezing followed by rapid freeze-substitution, as previously described (Kolotuev, 2014; Kolotuev et al., 2010). Dissected guts were immediately transferred to large high pressure freezing carriers filled with 20% bovine serum albumin for cryo-protection and frozen using the standard procedure according to the manufacturer’s instructions (High Pressure Freezing Machine HPF Compact 02, Engineering Office M. Wohlwend GmbH, Sennwald, Switzerland). Samples were substituted in an AFS2 machine (Leica) with 0.1% uranyl acetate diluted in anhydrous acetone and embedded in HM20 acrylic resin mix (Electron Microscopy Sciences). To assure precise orientation of the samples, the flat embedding procedure was used (Kolotuev, 2014).

### Electron microscopy image acquisition and analysis

Polymerized flat blocks were trimmed using a 90° diamond trim tool (Diatome, Biel, Switzerland) mounted on a Leica UC6 microtome. Transmission electron microscopy samples were analyzed with an FEI CM100 electron microscope operated at 80kV, equipped with a TVIPS camera piloted by the EMTVIPS program.

Samples for CLEM were sectioned at 100-150 nm thickness and transferred to wafers using an array tomography protocol (Burel et al., 2018; Kolotuev and Micheva, 2019). CLEM wafers were first imaged for fluorescence signal using a Zeiss fluorescent microscope equipped with DAPI and GFP filters using 20x and 60x objectives. To analyze the ultrastructure, sections on wafer were contrasted with uranyl acetate and lead citrate and observed using an FEI Quanta 250 FEG scanning electron microscope (FEI, Eindhoven). The imaging settings were as follows: accelerating voltage, 10kV; spot size, 5; image dimensions, 4096×4096; pixel dwell time, 10µs.

FIB-SEM tomography was done with a Helios 650 (FEI, Eindhoven). Fibbing conditions were 30 keV, 770 pA, 30-40 nm slice thickness (specified in text for each experiment) at a tilt angle of 52° and a working distance of 13 mm. For imaging the block face was tilted normal towards the electron beam (Kizilyaprak et al., 2015). The imaging conditions were: 2 keV, 800 pA, 20 µs dwell time, with a frame size of 6144 x 4096 and a pixel size of 9.7 mm. For publication, the image contrast was inverted.

IMOD (Kremer et al., 1996) was used to convert raw data from sequential sections to an MRC file stack and also used for alignment of serial sections and volumetric rendering. Adobe Photoshop was used for image adjustment, layers superposition, annotations, pseudo-coloring of image zones, and volume reconstructions.

### Volumetric rendering of FIB-SEM images

Serial sections were stacked and aligned using the cross-correlation function of IMOD, which was also used to trace and reconstruct specific regions. Drawing tools were used for outlining subcellular features (e.g., septate junctions, plasma membrane, nuclei, PAAC) on the EM layers. The 3D reconstruction surfaces were Meshed in Model View/ Objects tool. Images were captured using the Model View/Movie Montage tool and reformatted into .avi format using Fiji.

## Supporting information

Moreno-Roman_FigS1

Moreno-Roman_FigS2

Moreno-Roman_FigS3

Moreno-Roman_FigS4

Moreno-Roman_FigS5

Moreno-Roman_Video1

Moreno-Roman_Video2

Moreno-Roman_Video3

Moreno-Roman_Video4

Moreno-Roman_Video5

Moreno-Roman_Video6

## ACKNOWLEDGEMENTS

We are grateful to Allison Bardin, David Bilder, Nicolas Buchon, Joaquin de Navascues, Mikio Furuse, Yoshihiro Inoue, Henri Jasper, Sarah Siegrist, Norbert Perrimon, Nicolas Tapon, Charles Xu, and Drosophila stock centers (Bloomington *Drosophila* Stock Center (NIH P40OD018537), Vienna Drosophila Resource Center (Dietzl et al., 2007), Kyoto Drosophila Genomics and Genetic Resoruces)for fly stocks; Mikio Furuse, Claire Thomas, Steven Russell, and Xiaohang Yang for antibodies; and Jon Mulholland and Kitty Lee for microscopy support. Confocal microscopy was performed at the Stanford Beckman Cell Sciences Imaging Facility (NIH 1S10OD01058001A1, NIH 1S10OD010580). We thank David Bryant, Tobias Reiff, Daniel St. Johnston, Jia Chen, and members of the O’Brien lab for invaluable discussions. Fly extract was obtained from the *Drosophila* Genomics Resource Center (NIH 2P40OD010949).

## FIGURES and CAPTIONS

**Figure S1.**
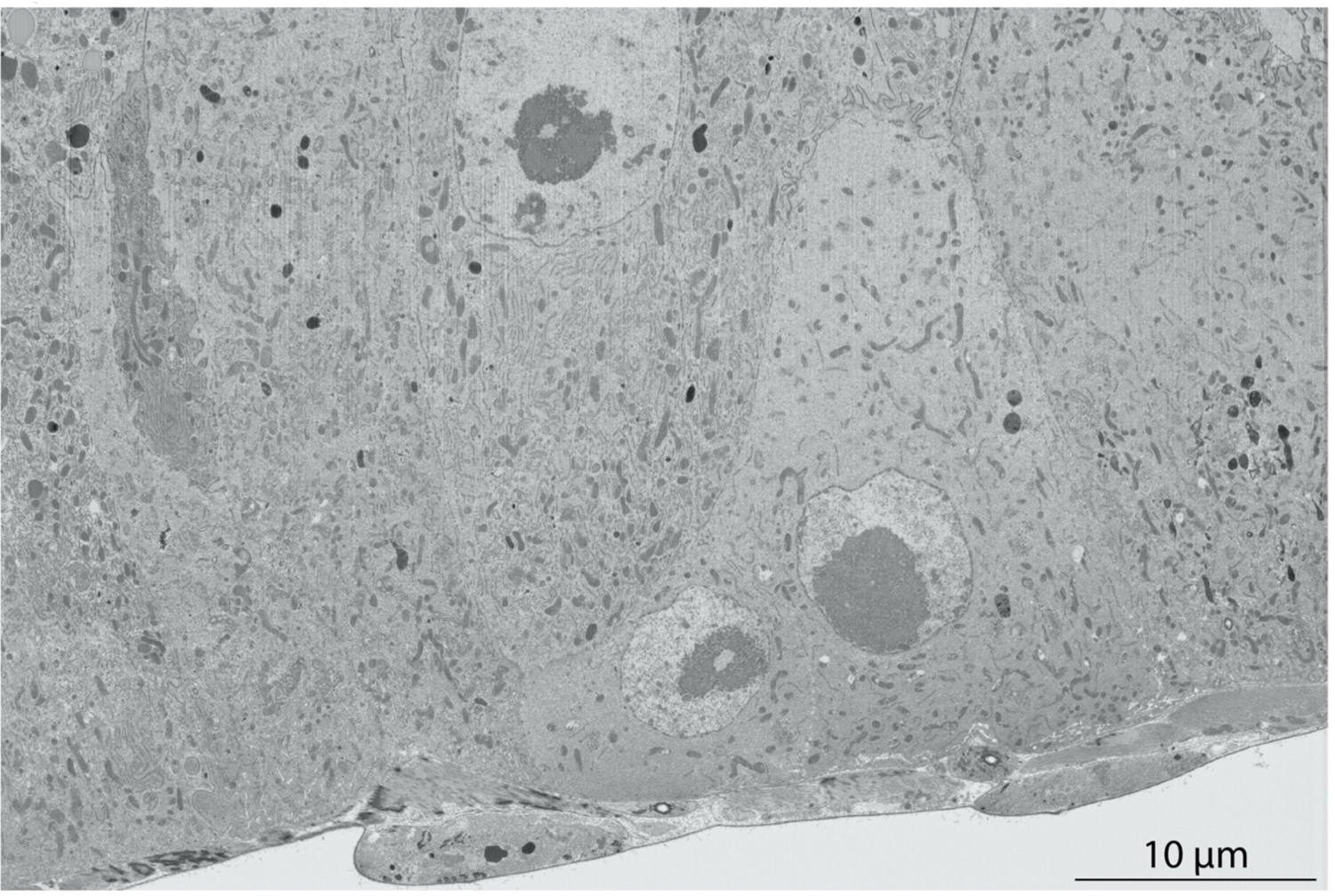
(related to Figure 2D). High resolution view of FIB-SEM section shown in Figure 2D. 30 nm-thick sections were cut with a gallium ion beam at 30 keV and 770 pA. Images were taken with the electron beam at 2 keV, 0.8 nA, 2 mm working distance, 20 µs dwell time, 6144×4096 pixel frame size. Pixel size 9.7nm. Full genotypes in Table 1.

**Video 1 (related to Figure 2C). Three-dimensional ultrastructure of nascent SJ between Su(H)-GFP::nls^+^ enteroblast and mature enterocytes.**

Tomographic reconstruction of 30 serial images, including the image in Fig. 2B, from midgut expressing *Su(H)-GFP::nls.* Serial sections were cut with a gallium ion beam at 10kV, spot size 5, pixel frame size 4096×4096, pixel dwell time 10µs. Pixel size 8.7nm. Slice thickness, 150nm. Volume of reconstruction, 35.6um x 35.6um x 4.5µm. Full genotypes in Table 1.

**Video 2 (related to Figure 2E). Three-dimensional ultrastructure of SJ ‘cap’ between enteroblast and mature cells.**

Tomographic reconstruction of 413 serial FIB-SEM images, including the image shown in Figs. 2C and S1. Volume of reconstruction, 55 µm x 36.6 µm x 12.3 µm. Slice thickness, 30nm. Full genotypes in Table 1.

**Figure S2.**
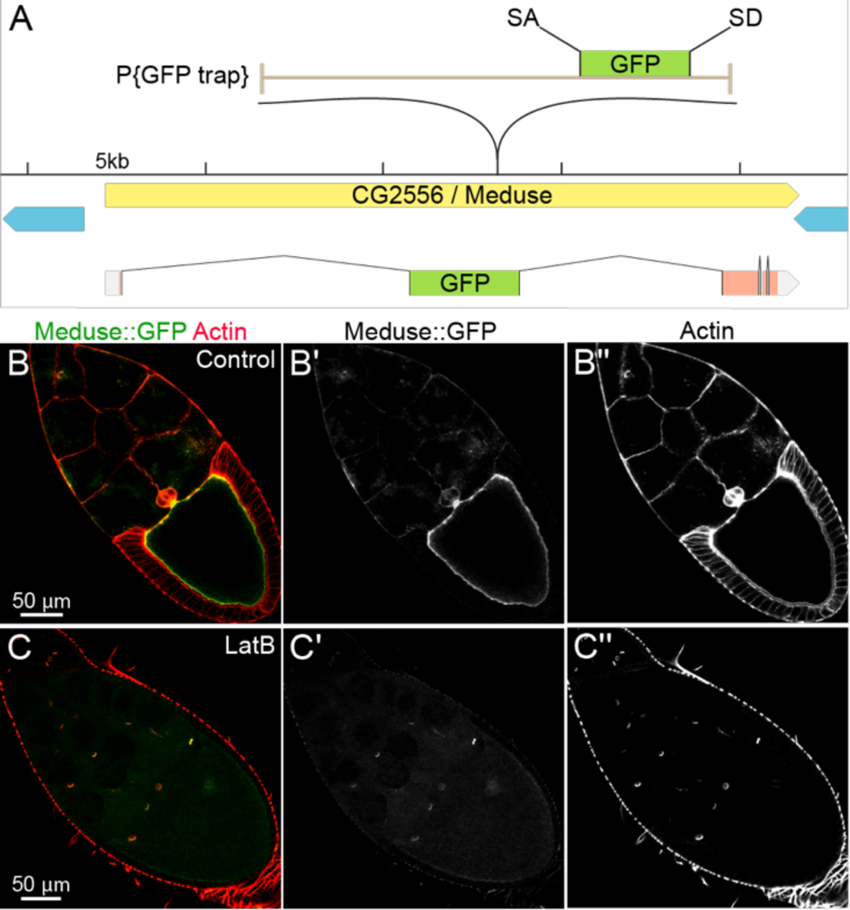
(related to Figure 3A-C). The A142 splice trap transposon is inserted into CG2556/Meduse, a novel protein that co-localizes with actin filaments. (A) Genomic location of the splice trap transposon in the A142 line. The insertion was mapped by inverse PCR and genomic PCR to the large first intron of CG2556, approximately 10.6 kb downstream of the splice site in Exon 1. The transposon is inserted in the proper orientation to capture transcripts from CG2556, which would result in an N-terminal GFP tag on the nearly undisrupted protein (Exon 1 encodes only 7 amino acids including the initiator Met). The tentacular appearance of the fusion protein in oocytes prompted us to name the gene Meduse (Mdu). (B) Mdu::GFP co-localizes with cortical actin filaments in Stage 10 oocytes. (C) Latrunculin B (LatB) treatment disrupts cortical actin filaments (red, Rhodamin-phalloidin) in the oocyte and leads to abrogation of the oocyte Mdu::GFP signal. Note that LatB does not disrupt actin in ring canals; localization of Mdu::GFP to ring canals is visible in Panels C and C’. Full genotype in Table 1.

**Figure S3.**
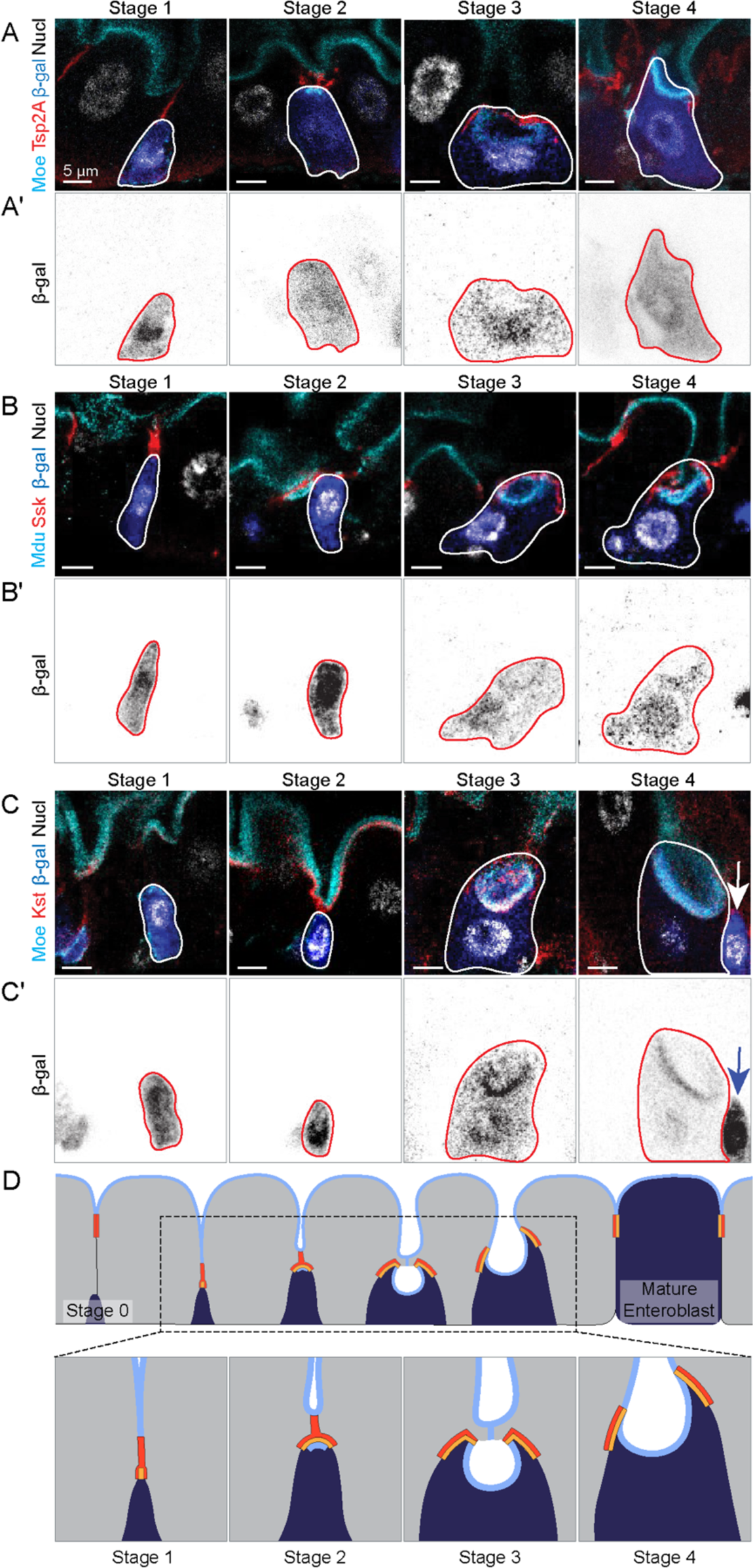
(related to Figure 3A-C). Immunostaining of *Su(H)*-driven β-galactosidase in integrating cells. (A-C) The same four-channel images shown in Figure 3 are repeated above the corresponding, single-channel images of β-galactosidase immunostain. The presence of β-galactosidase in Stage 3 and Stage 4 cells demonstrates that these cells derived recently from enteroblasts. During acquisition of the Stage 3 and 4 images, the gain was increased compared to Stages 1 and 2 to visualize lower levels of β-galactosidase. Arrowheads in C and C’ point to a Stage 1 enteroblast next to the Stage 4 pre-enterocyte; at the higher gain necessary to visualize β-galactosidase in the Stage 4 pre-enterocyte, β-galactosidase intensity in the Stage 1 enteroblast is overexposed. Images are projections of short confocal stacks. (D) Four-stage model of epithelial integration. A newborn enteroblast (Stage 0) forms SJ contacts between their apex and the basal edge of enterocyte-enterocyte SJs (Stage 1). The enteroblast-enterocyte SJ broadens, and apical markers accumulate at its cytoplasmic face (Stage 2). The enteroblast grows into a pre-enterocyte, characterized by the expansion of apical markers into a PAAC and broadening of the SJ into a diaphragm-like sheet (Stage 3). The PAAC’s lumen fuses with the gut’s central lumen, and SJs remodel to circumscribe the cell (Stage 4). Eventually, the concave lumenal-apical surface everts to form the convex apical surface that characterizes mature enterocytes. Enteroblast/pre-enterocyte SJs shown in orange, mature enterocyte SJs in red, and apical surface in light blue. Full genotypes in Table 1.

**Video 3 (related to Figure 3A). 360° confocal reconstruction of a Stage 3 pre-enterocyte shows that the PAAC’s apical membrane is distinct from the gut’s lumenal-apical surface.**

Video shows reconstructed 360° view of a Stage 3 pre-enterocyte, labeled by Su(H)-driven β-galactosidase. The pre-enterocyte is surrounded by two mature enterocytes, and a pair of small, basal progenitor cells is visible between the pre-enterocyte and one of the mature enterocytes. The apical marker Moesin::GFP outlines the lumenal-apical surface of the mature enterocytes, the PAAC in the pre-enterocyte, and the entire cortex of the progenitor cells. The SJ protein Tetraspanin2A forms a convex web that covers the apex of the pre-enterocyte. Nuclei are labelled with DAPI. Full genotype in Table 1.

**Video 4 (related to Figure 4C and 4D). 7.5-hour continuous time-lapse of SJ dynamics during enteroblast-enterocyte differentiation.** Windowmount imaging of midgut expressing *GS5961-tsp2A::GFP* and *Su(H)-mCherry*. Planar (top) and ortho (bottom) views of the same tissue volume are shown. In ortho view, the dotted white line indicates the basal surface of the differentiating cell. Arrowhead points to SJs (Tsp2A::GFP) associated with the differentiating, mCherry^+^ cell analyzed in Fig. 4B. Dynamics of the SJ in ortho view are consistent with the four-stage mechanism inferred from Fig. 3: Nascent contact (Stage 1), broadening and expansion over the cell apex (Stages 2-3), and central hollowing and lateral extension (Stage 4). Full genotype in Table 1.

**Video 5 (related to Figure 4E). 3.75-hour continuous time-lapse movie of PAAC development.** Windowmount imaging of midgut expressing *moesin::GFP* and *ubi-his2av::RFP*. The Moesin::GFP channel (magma LUT) is shown without (left) and with (right) the His2av::RFP channel (grayscale). At 30 min, a faint Moesin::GFP-labelled structure (arrowhead) forms at the basal side of the gut’s lumenal-apical surface. The concave shape of this structure is similar to PAACs in fixed samples (Fig. 3, Stage 3; Video 3). From 30-225 min, the putative PAAC deepens, and its GFP-labelled boundary brightens and thickens. Full genotype in Table 1.

**Figure S4.**
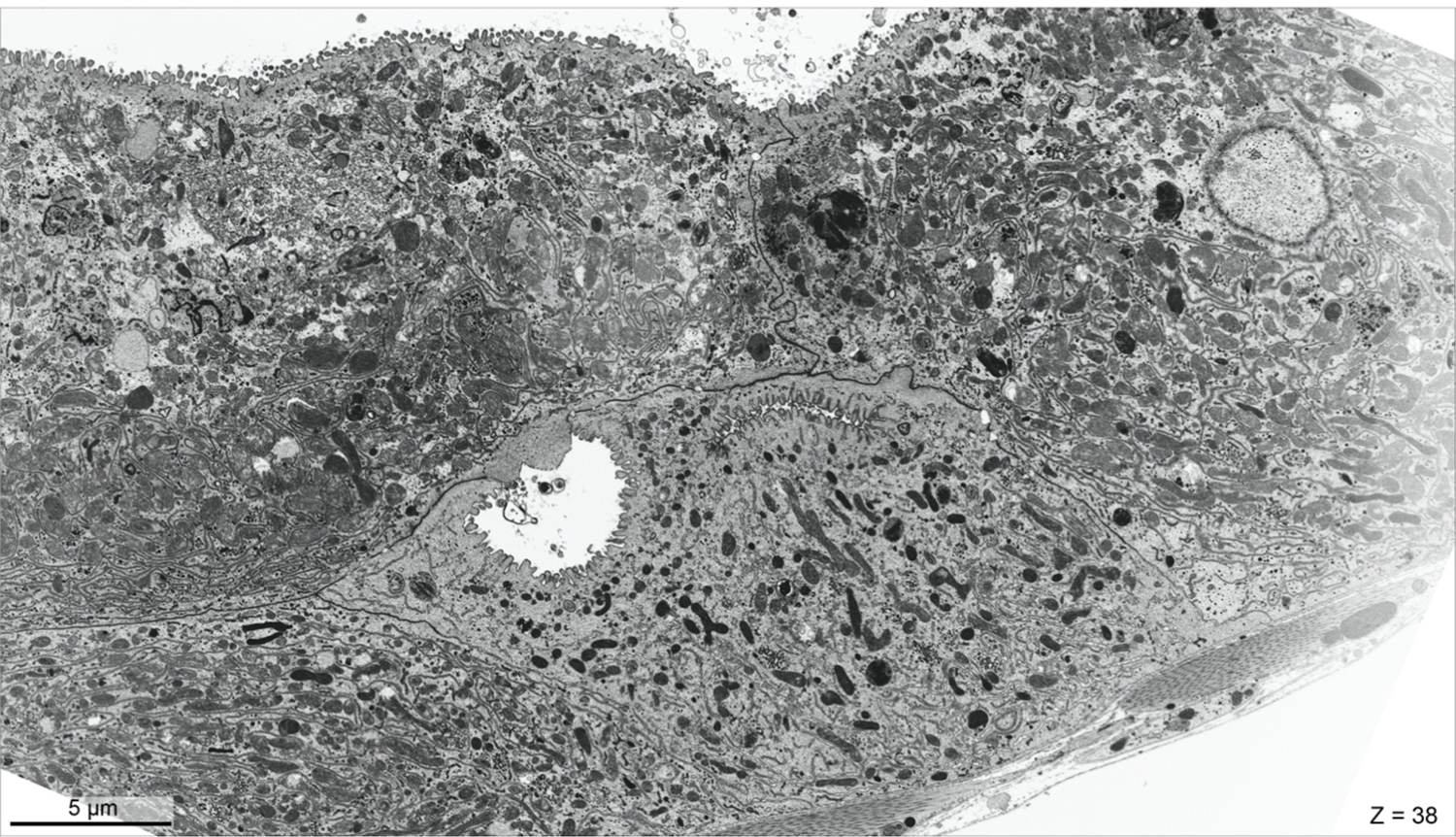
(related to Figure 5A). High resolution view of FIB-SEM section shown in Figure 4A. 40nm-thick sections were cut with a gallium ion beam at 2kV, 0.8 nA, 4.2mm working distance, 5µs dwell time, 6144×4096 frame size. Pixel size 9.7nm. Full genotype in Table 1.

**Video 6 (related to Figure 5A-D). Three-dimensional ultrastructure of PAAC, PAAC precursor, and their associated pre-enterocyte.** Tomographic reconstruction of 200 serial FIB-SEM images, including a cropped version of the image shown in Figures 5A and S4. 360° rotation reveals the ellipsoid and allantoid shapes of the PAAC and PAAC precursor, respectively, and also reveals holes in the SJ in which the pre-enterocyte and enterocyte membranes have separated to form the intercellular lumens. Volume of reconstruction: 40.2 µm x 23.9 µm x 8 µm. Slice thickness, 40 nm. Full genotype in Table 1.

**Figure S5.**
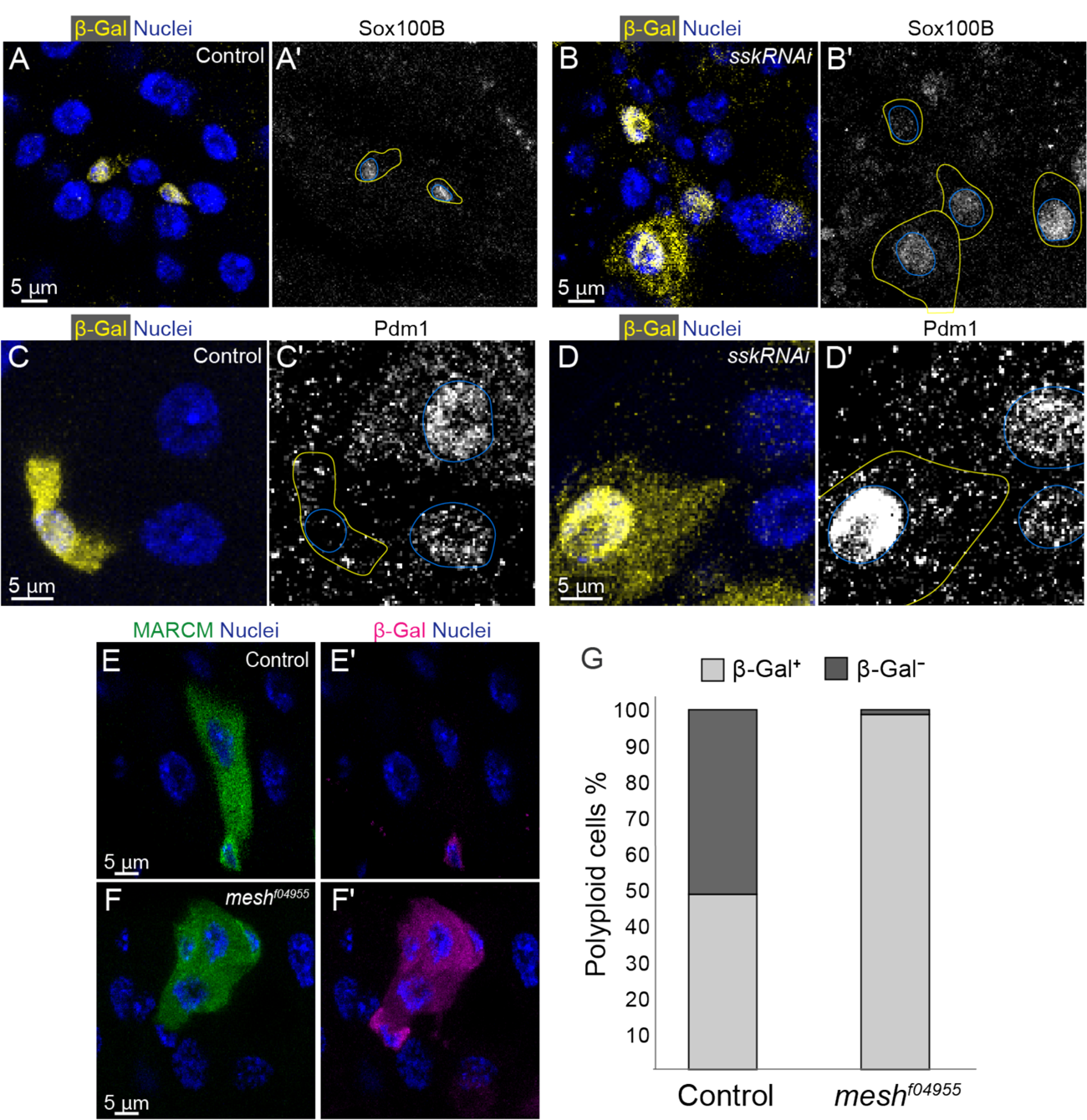
(Related to Figure 6). SJ-less cells exhibit an abnormal mix of enteroblast and enterocyte features. (A-B) *Su(H)^ts^>sskRNAi* cells express the stem cell/enteroblast transcription factor Sox100B. Planar views of control *Su(H)* cells (A) and *Su(H)^ts^>sskRNAi* cells. (B). Left panels: Yellow, *Su(H)-lacZ* (β-gal immunostain); blue, nuclei (DAPI). Right panels: Sox100B immunostain. Yellow and blue outlines in right panels indicate *Su(H)-lacZ^+^* cells and their nuclei, respectively. (C-D) *Su(H)^ts^>sskRNAi* cells express the enterocyte-specific transcription factor Pdm1. Control *Su(H)* cells (C) and *Su(H)^ts^>sskRNAi* cells are shown. Left panels: Yellow, Su(H)-lacZ (β-gal immunostain); blue, nuclei (DAPI). Right panels: Pdm1 immunostain. Yellow and blue outlines in right panels indicate *Su(H)-lacZ^+^* cells and their nuclei, respectively. (E-G) *Su(H)-lacZ* expression in stem cell (MARCM) clones that are either control (E) or *mesh*-null (*mesh^f04955^*) (E). 4-day clones are labelled with GFP (green). β-gal immunostain (red) identifies *Su(H)-lacZ*-expressing cells. Nuclei are stained with DAPI (blue). In control clones, 52% of polyploid cells have lost β-Gal staining (n=39 polyploid cells from 133 clones, N=3 midguts), implying that terminal differentiation of polyploid cells to enterocyte fate is complete. In *mesh*-null clones, only 1.1% of cells has lost β-gal staining, implying that nearly all cells, despite being polyploid, have not completed terminal differentiation (n=89 polploid cells from 149 clones, N=3 midguts). All scale bars, 5 µm. Full genotypes in Table 1.

