## Supplementary figures and images for "Progenitor cell integration into a barrier epithelium during adult organ turnover"

### Moreno-Roman_FigS1

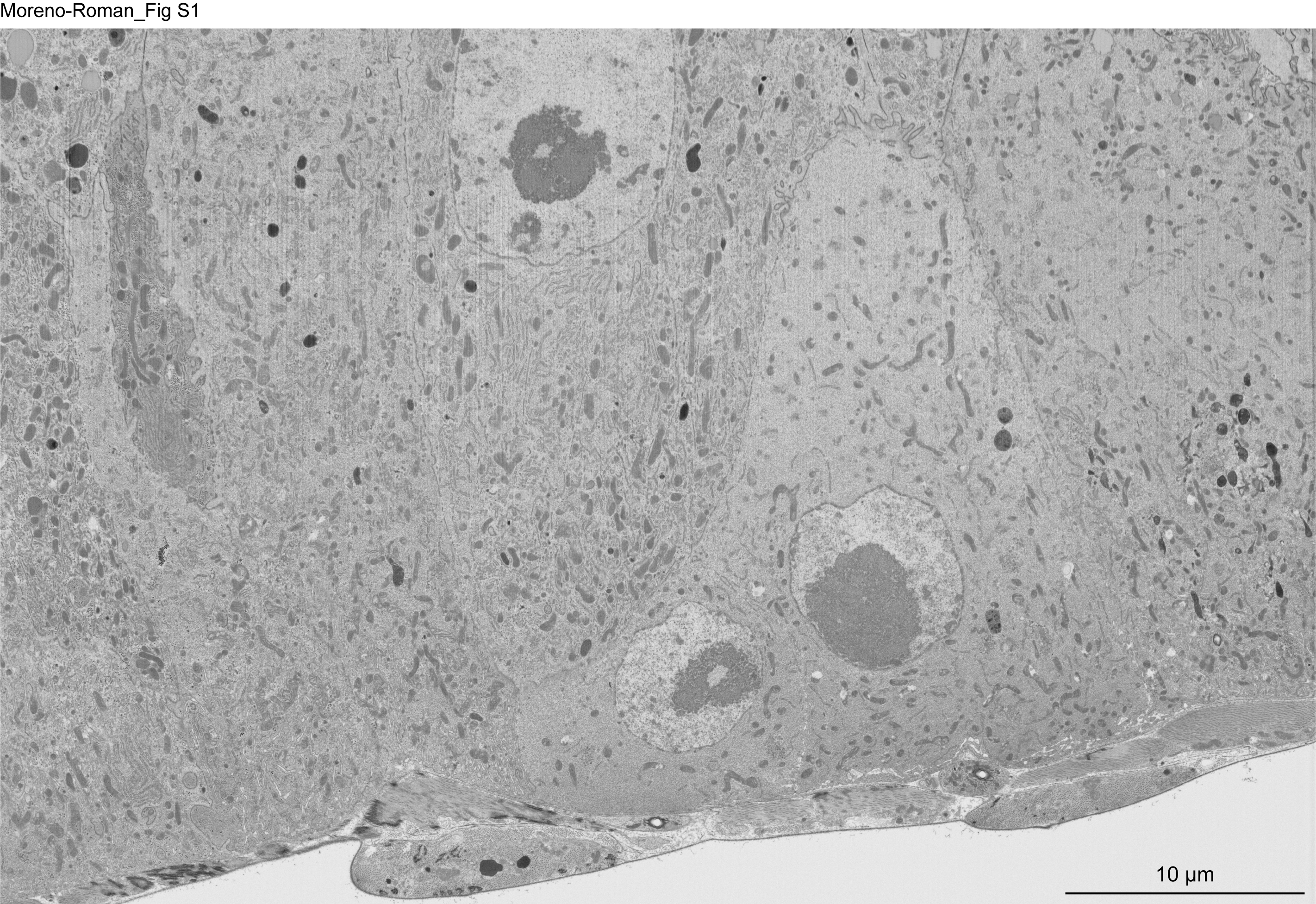

### Moreno-Roman_FigS2

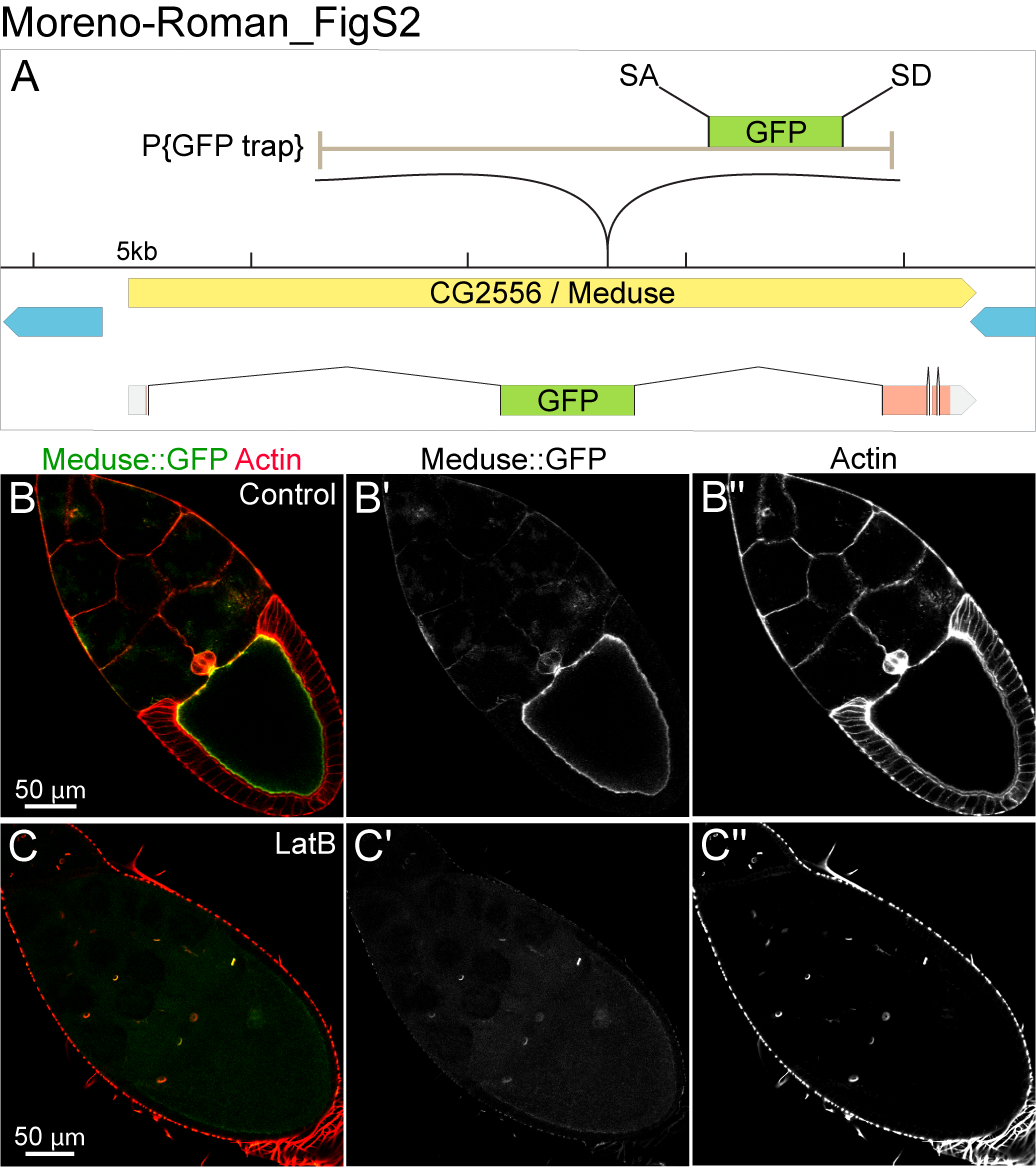

### Moreno-Roman_FigS3

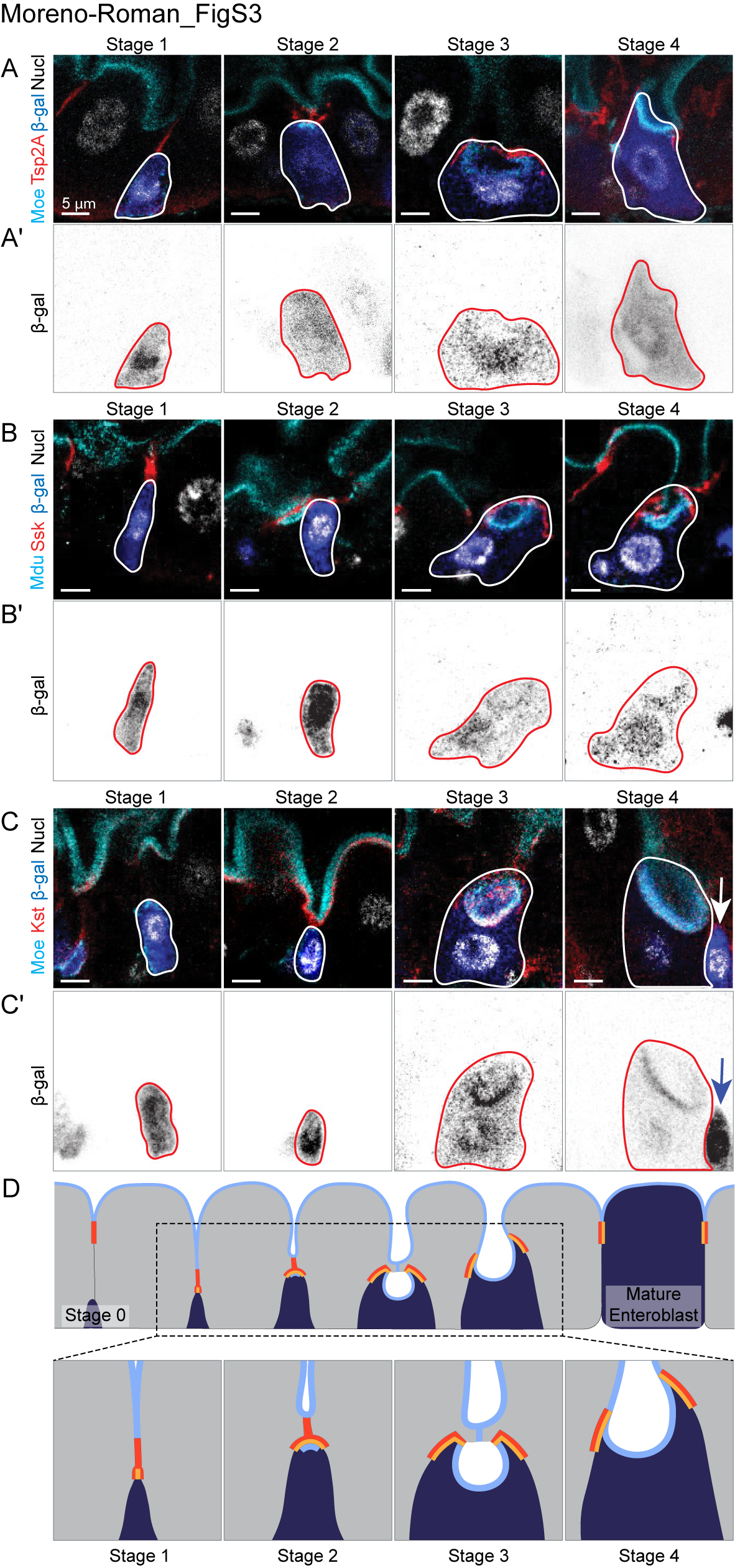

### Moreno-Roman_FigS4

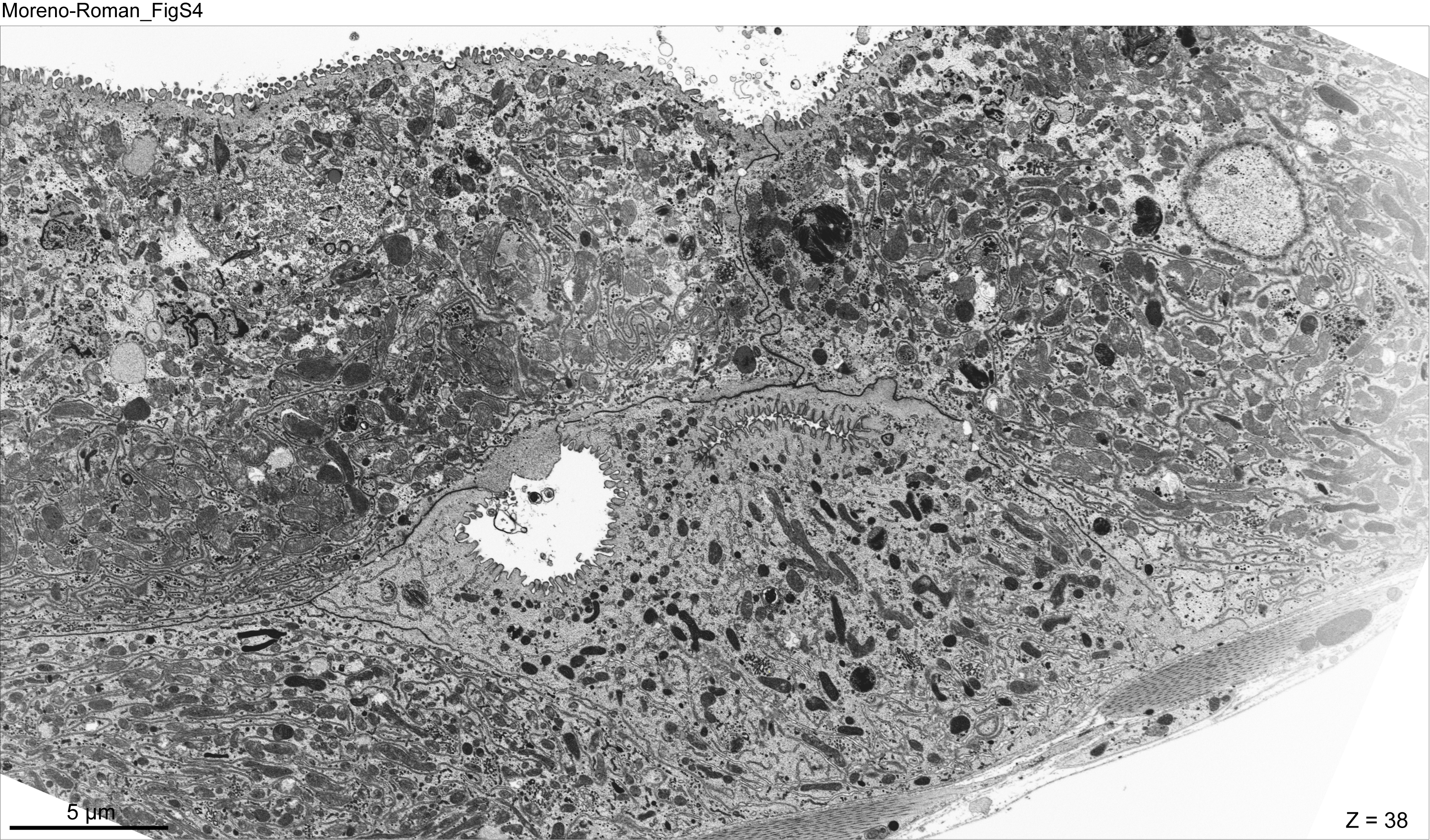

### Moreno-Roman_FigS5

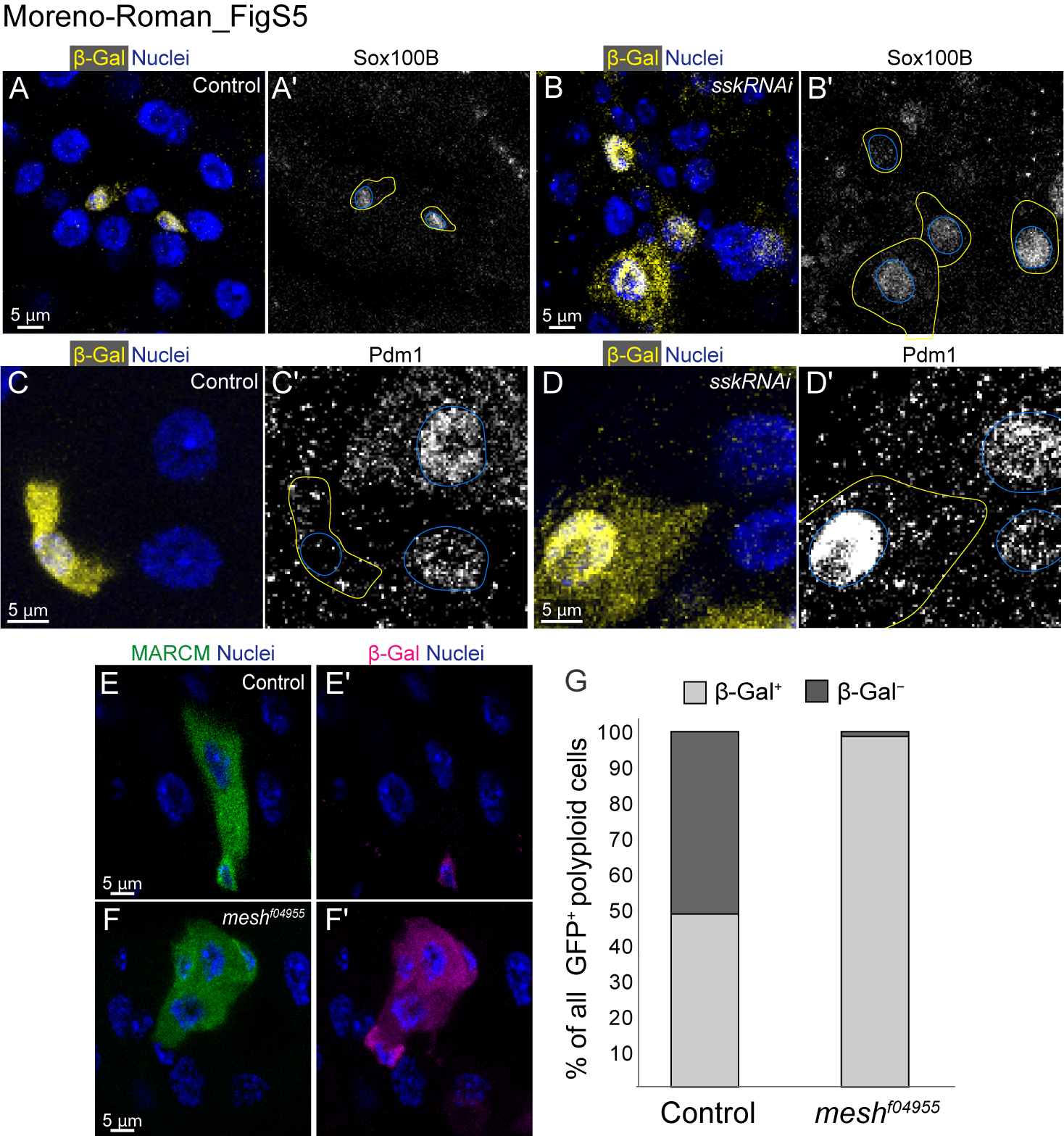
